# Short-term gonadal cultures are sufficient for germline transmission in a songbird

**DOI:** 10.1101/2025.04.25.650428

**Authors:** Matthew Biegler, Elijah Harter, Asha V. Sidhu, Christina Szialta, Gillian Durham, Lotem Tchernichovski, Paul Collier, Ji-Dung Luo, Wei Wang, Ryan MacIsaac, Kirubel Belay, Thomas Carroll, Anna L. Keyte, Erich D. Jarvis

**Author notes:** Co-second authors (last name alphabetical) (Elijah Harter, Asha V. Sidhu, Christina Szialta).

## Abstract

1

Primordial germ cells (PGCs) are germline stem cells that develop into sperm or egg cells and are valuable for avian biobanking and the propagation of donor-derived offspring. However, in non-poultry birds the long-term maintenance and self-renewal of PGCs *in vitro* remains challenging. This limitation hinders biobanking in other avian clades, particularly in the zebra finch and other songbirds that uniquely possess a germline restricted chromosome (GRC). Here, we generated and compared short-term cultures of chicken and zebra finch PGCs from the embryonic gonads or blood, as well as established long-term cultures of chicken PGCs. Using single-cell RNA sequencing, we found that the transcriptome profile of long-term chicken gonadal cultures were exclusively PGCs, whereas the short-term chicken and zebra finch cultures represented a heterogeneous mixture of cell types. The zebra finch culture further included rapidly differentiating PGCs, as well as a germ cell type not previously identified in the embryonic songbird gonad. Although zebra finch short-term gonadal cultures did not yield robust long-term PGC cultures, short-term cultured PGCs were able to integrate into host zebra finch gonads after injection into the dorsal aorta, contribute to gametic populations in adult chimeras, and give rise to phenotypically- and genomically-validated offspring. This study provides a foundation for using short-term gonadal cultures to derive donor and transgenic offspring in songbirds and further explore the unique developmental genetics of PGCs across the avian clade.

**Summary:** Beyond poultry, the long-term culture of self-renewing primordial germ cells (PGCs) remains a challenge. Here, we compare the cell population heterogeneity and reproductive viability of gonadal cultures for the zebra finch, a songbird model of vocal learning, with established chicken PGC protocols. Using single-cell RNA sequencing, we identify the rapid differentiation of zebra finch gonadal germ cells *in vitro*, including germline identities not previously noted in the embryonic gonad. In comparison, these differentiated cell profiles were also found in zebra finch blood PGC culture conditions, but not identified in short- or long-term chicken PGC cultures. Host embryo injections of these short-term zebra finch gonadal cultures resulted in germline chimeric animals, but at lower rates of gonadal reconstitution compared to chicken. Nonetheless, these cultures allowed for the derivation of zebra finch germline chimeras that yield phenotypically- and genomically-validated offspring from cultured PGCs.

## 2 Introduction

The isolation and *in vitro* propagation of avian primordial germ cells (PGCs), a type of germline stem cell that gives rise to oogonial and spermatogonial cells, provides a powerful resource for creating germline transgenic animals. Methods using PGCs are important in clades where zygotic manipulation is difficult, such as in birds^1,2^. Avian PGCs take a unique vascular migration to the gonads^3–5^. In the chicken, PGC cultures obtained from either HH14-16 blood^6–8^ or HH28^9–11^ gonad has provided reproductively-viable material for surrogacy and biobanking. These methods have been successfully implemented for multiple breeds^12–14^, enabling the maintenance of genetically modified or heritage breed chickens through PGC cells lines that do not require active breeding stock or cellular reprogramming.

Other than in chicken, and recently in goose^15^, the robust maintenance and self-renewal of PGCs in other avian species has not been obtained beyond short-term blood or gonadal PGC cultures that persist from a few days to several weeks^16–19^. This lack of suitable culture methodology represents a significant gap in the feasibility of using avian PGCs for germline transmission and biobanking. In songbirds, PGC culture systems are further complicated by the presence of a germline-restricted chromosome (GRC), which presumably has a role in germline development^20,21^. Therefore, while other avian species may eventually benefit from embryonic stem cell culture and somatic cell reprogramming^22,23^, germ cells are likely the only option to comprehensively biobank the songbird genome and derive gene edited offspring.

In the zebra finch, a popular songbird model for vocal learning research, germline transmission has been recently demonstrated using zebra finch blood PGC cultures cultured for up to two weeks^24^. These culture conditions allowed for lentiviral transduction prior to blastodermal injection, which yielded transgenic offspring from germline chimeras. However, the use of gonadal cultures offers an important alternative to the technically demanding blood PGC collection protocol^25^, particularly when the samples can only be collected opportunistically. Gonadal PGC culture protocols in chicken were also adapted for the zebra finch, enabling proliferation for up to four weeks, which allowed for manipulation and gonadal reconstitution^26–28^. To date, however, no demonstration of germline transmission using zebra finch gonadal cultures has been reported.

One issue in zebra finch gonadal cultures may be germline heterogeneity and differentiation. Our recent study^29^ comparing single-cell RNA sequencing (scRNAseq) datasets of the developing zebra finch and chicken HH28 gonad found that both male and female zebra finch PGCs undergo *FOXL2L*-associated germ cell differentiation as early as HH25. These differentiated germ cells are highly proliferative and lack migratory PGC markers (e.g., *NANOG*, *PRDM14*). As this developmental process does not occur in chicken male embryos and only begins in females around HH36^30,31^, we wondered how this species difference might influence zebra finch PGC cultures *in vivo*.

In this study, we sought to characterize the molecular differences between *in vitro* chicken and zebra finch PGC cultures, determine the proportion that are likely viable for germline transmission, and exploring the use of gonadal zebra finch PGC cultures as an alternative to derive offspring. We identify key species differences in the proportion of PGCs expressing pluripotent and migratory markers over time. Despite extensive germline differentiation *in vitro,* deriving donor-derived zebra finch offspring from embryonic gonads was possible. This work highlights the viability of multiple songbird tissue types for PGC collection and offers a foundation to expand avian germline transmission methods beyond the chicken.

## 3 Results

### 3.1 Zebra finch gonadal PGC cultures undergo rapid differentiation

Replicating findings with our prior media conditions^26^, short-term pooled male and female zebra finch gonadal cultures (3-6 days *in vitro* (DIV)) consisted of a non-adherent fraction of cells with round morphology and light reflectance characteristic of PGCs, and an adherent fraction of somatic gonadal cells^26^ (**Figure S1A**). As noted in previous studies^24^, the non-adherent zebra finch PGCs generally presented a more clustered morphology in suspension than in the chicken (**Figures S1B** and **S1C**). We separated these fractions from two distinct DIV3 pools one DIV6 pool and enriched for live cells by FACS-based dead cell exclusion with DAPI. We then performed scRNAseq on the non-adherent and the adherent fractions in parallel, to prevent batch effect differences (**Figure S2A**). Live, singlet sorted cells comprised ∼60-80% of the total non-adherent population (**Table S1**).

#### 3.1.1 Non-adherent zebra finch gonadal culture fraction

Quality filtering of the non-adherent datasets yielded 13,785 high-quality cell barcodes (**Figures S2A**-**B**; **Table S2**). Cell type assessment, using canonical marker genes and label transfer analysis, identified most cells as germ cells across each sample (**Figure 1A**). Cluster analysis of the integrated DIV3 and DIV6 datasets yielded discrete gene marker profiles (**Figures 1B** and **S2C**). Overall, we identified four germ cell clusters (c1, c2, c3, and c5.1; 11,378 cells), two immune clusters (c4 and c7; 2,209 cells), and three small clusters of other somatic cell types (c5.2, c6, and c8; 260 cells). Despite cell cycle normalization, we noted distinct cell cycle phase representation across clusters, with cluster c2 predominantly containing G2M and S phase cells (**Figure S2D**).

**Figure 1.**
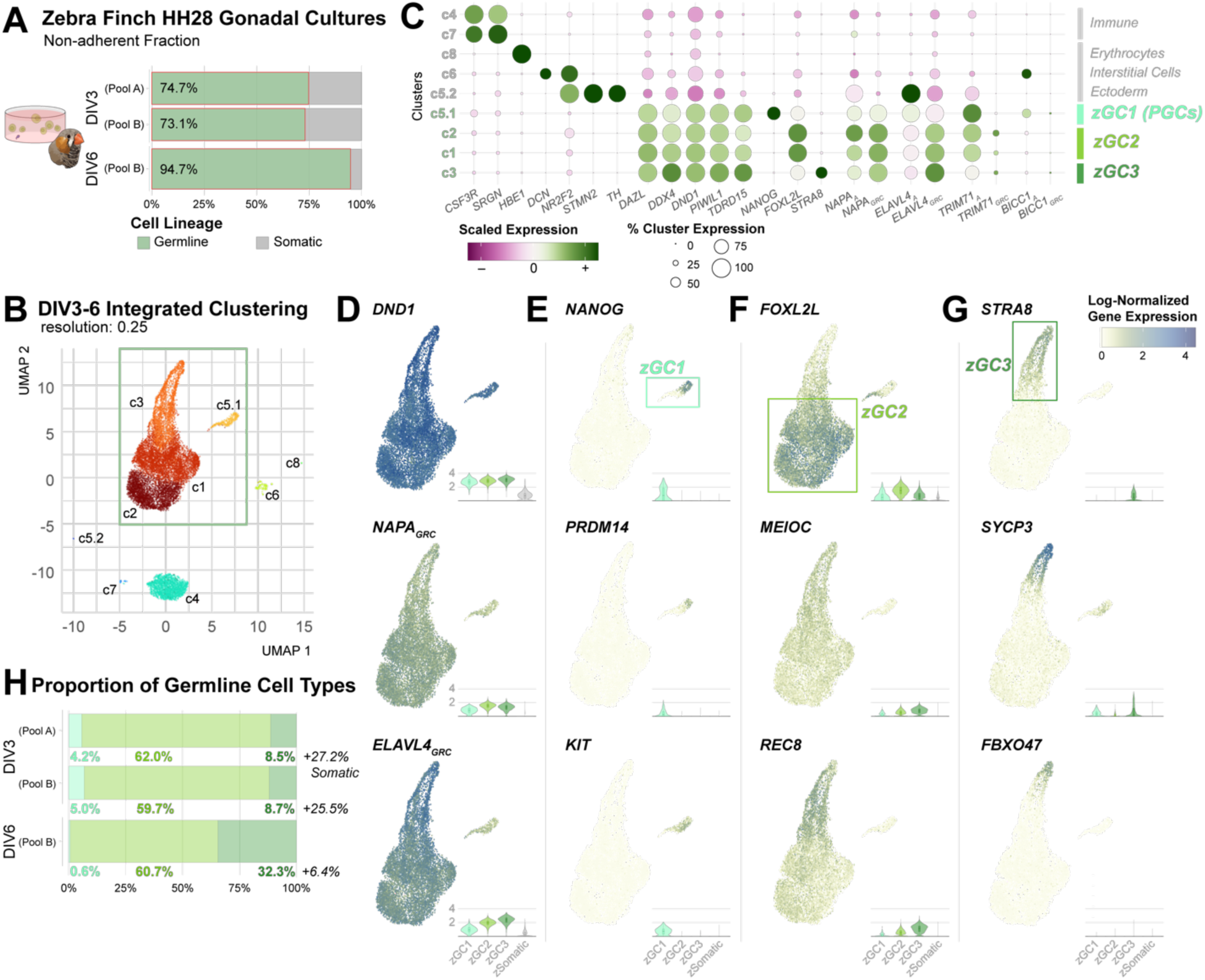
scRNAseq profiling of non-adherent cell types from short-term zebra finch gonadal cultures. (**A**) Relative proportion of cell types determined from scRNAseq analysis among the non-adherent cells in DIV3 and DIV6 cultures from zebra finch embryonic gonads. (**B**) UMAP dimensional reduction of integrated non-adherent zebra finch gonadal cell cultures, colored and labeled by nearest-neighbor assigned cluster. (**C**) Dotplot of select gene markers expressed in each cluster that define specific cell types. (**D**) Log-normalized gene expression overlaid on a UMAP of the integrated germline subset and as a violin plot for each germ cell type, highlighting broad germline markers; (**E**) zGC1 (PGC) enriched markers; (**F**) zGC2 enriched markers; and (**G**) zGC3 enriched markers. (**H**) Relative proportion of germ cell types determined from scRNAseq analysis among the non-adherent cells in DIV3 and DIV6 cultures from zebra finch embryonic gonads.

We validated the four germline clusters with highly expressed canonical markers like *DAZL*, *DDX4*, and *DND1* (**Figure 1C**). We noted low-level expression of these markers in a high proportion of somatic clusters, indicative of ambient RNA despite cell sorting and empty droplet RNA recalibration. Other germline markers with lower expression levels (e.g., *TDRD15*) or specific germ cell types (e.g., *NANOG*) were more strictly expressed (**Figure 1C**). This non-specific signal may be due to several factors, including high post-sort cell mortality, the inclusion of DAPI-negative cell debris, and the adherence of cell debris to live cells.

When we compared the clusters with *in vivo* cell types defined in our HH28 zebra finch gonadal scRNAseq dataset^29^, and we found most *in vitro* clusters discretely corresponded to *in vivo* germline or somatic cell types (**Figure S3A**). Cluster c5.1 mapped to zebra finch zGC1 cells (considered gonadal PGCs), while clusters c1-3 mapped primarily to zGC2 cells (considered differentiated gonadal germ cells). An assessment of sex chromosome gene expression identified both male and female cells in each germ cell cluster (**Figure S2D** and **S2E**), consistent with the sexually monomorphic germline differentiation patterns seen in the HH28 zebra finch gonad^29^. Cluster c6 (174 cells) mapped back to multiple somatic cell types, including intermediate mesodermal (IM) progenitor and Interstitial cell subtypes.

We also assessed germline expression of several genes present on the germline restricted chromosome (GRC)^32^. The GRC is unique genomic element in songbirds^33^ (and no other avian clade), which undergoes programmed DNA elimination from the somatic lineage early in embryonic development^34^. Compared to robust *DND1* expression across all germline clusters (**Figure 1D**), both *NAPA*_GRC_ and *ELAVL4_GRC_* were more variable across clusters. Both GRC genes were downregulated in the cluster c5.1, while cluster c3 had higher expression of *ELAVL4_GRC_* and lower expression of *NAPA*_GRC_ compared to c1 and c2.

We then compared differentially expressed genes (DEGs) across all clusters (**Table S3**). Cluster c5.1 was positively marked by 4,438 genes, including several canonical PGC markers that were not present in other germ cell clusters (*NANOG*, *PRDM14, KIT*; **Figure 1E**). In contrast, clusters c1-3 were most closely aligned with *in vivo* zGC2 cells expressed the differentiation markers *FOXL2L, MEIOC,* and/or *REC8* (**Figure 1F**). Combined, clusters c1 and c2 were positively marked by 2,440 unique DEGs (**Table S3**), while 1,169 DEGs were identified between them, mostly differing by dosage and proportion (**Table S4**). Interestingly, cluster c3 had decreased *FOXL2L* expression compared to clusters c1 and c2, as well as an increase in *MEIOC* and *REC8* (**Figures 1F**). Cluster c3 had upregulation of 3,014 DEGs (**Table S3**), many of which were not highly expressed *in vivo* at HH28, including *STRA8* and *SYCP3* (**Figure 1G**), with the latter showing modest expression in the zGC1 population as well. Several cluster c3 DEGs, such as *FBXO47*, were only identified in a small subpopulation of this cluster (**Figure 1G**). Accordingly, we designated cluster c3 as a third germ cell type, zGC3, transitioning from the zGC2 clusters.

Finally, we assessed whether there were proportional changes in germ cell cluster size between samples and timepoints. Between the DIV3 batches, consistent proportions of the germline clusters were identified (**Figures 1H**). In contrast, between DIV3 and DIV6 batches, the proportion of PGCs (cluster c5.1) decreased from 4.2 and 5% to only 0.6% (**Figure 1H**), while the proportion of zGC3 cells increased four-fold. Overall, the zebra finch non-adherent cell demographics show that PGCs are present early on but proportionally decline with time with concomitant increase in more differentiated germ cells beyond what is found at HH28.

#### 3.1.2 Adherent zebra finch gonadal culture fraction

We next sought to understand the representation of the adherent layer in the short-term cultures. We generated two scRNAseq datasets of the adherent cell layer of DIV3 and DIV6 cultures, derived from the same Pool B samples used for the non-adherent cells. Quality-filtering of the adherent datasets yielded 7,024 cells that resolved into seven clusters (**Figure S4A** and **Table S2**). Gene marker analysis (**Table S5**) demonstrated that the adherent layer was predominantly composed of four interstitial cell clusters (**Figures S4B-D**), while other small clusters were identified as containing germline (c5; 132 cells), immune (c6, 51 cells), or ectoderm (c7; 15 cells) populations. Within the interstitial clusters, some differed in the inferred proportion of male (c1, c3) or female (c2, c4) cells (**Figure S4C**). Reference mapping of the adherent cells to zebra finch *in vivo* HH28 gonadal scRNAseq datasets^29^ further validated that clusters c1-c4 were most like Interstitial and IM Progenitor populations, while cluster c5 mapped to germ cells and c6 to immune cells (**Figure S4E**). The above findings in both non-adherent and adherent cells indicate that the cell culture types at these early time points of DIV3 mostly maintain their gonadal molecular properties, with some more differentiation by DIV6.

To rule out distinct or differentiating germline populations between the culture fractions, we assessed the identity of cells within the germline cluster. Germ cells in zebra finch non-adherent cluster c5 were predominantly *DAZL*/*FOXL2L+* (**Figure S4F**), mirroring the overwhelming zGC2 representation in the non-adherent layer. Relatively smaller numbers of *NANOG*+ and *STRA8*+ cells were identified in the DIV3 and DIV6 zebra finch germ cell clusters (**Figure S4F**), which further paralleled the non-adherent population demographics and did not suggest a significant distinct germline population in the adherent layer.

### 3.2 Chicken gonadal culture conditions maintain PGC identity

We next compared short-term zebra finch gonadal culture demographics to both short-term (**Figure S1B**) and long-term (**Figure S1C**) chicken gonadal culture demographics. Like for the zebra finch, we dissociated pools of male and female HH28 gonads and cultured them using previously reported high-serum chicken PGC media conditions^11,12,35^. Only DAPI-negative singlet cells were retained by FACS in both the adherent non-adherent fractions for scRNAseq library preparation. The live singlet FACS sorted cells comprised ∼15-60% of the total non-adherent population, lower than the ∼60-80% for the zebra finch (**Table S1**), and may indicate species-specific sensitivities of gonadal cultures to dissociation and handling. From these cultures, we generated four scRNAseq chicken gonadal culture datasets: three non-adherent datasets from short-term DIV3-DIV6, and one adherent dataset from DIV3. We also generated a scRNAseq dataset from a DIV150 male gonadal PGC line grown on a mitomycin-treated mouse embryonic fibroblast (MEFs) feeder layer, derived from a single embryo short-term gonadal culture (**Table S1**).

#### 3.2.1 Non-adherent chicken gonadal culture fraction

Quality filtering of scRNAseq datasets for the three non-adherent, short-term chicken gonadal cultures yielded 4,210 high-quality cells (**Table S2**). In the short-term cultures, marker-based cell type inference identified a much lower proportion of *DDX4*/*DAZL*/*DND1*+ germ cells compared to the zebra finch non-adherent samples (**Figure 2A** vs **1A**), a parallel to the proportional germ cell differences in the HH28 gonad of each species^26,36^. Clustering of the combined short-term data segregated the cells into 11 nearest-neighbor clusters, of which four were germ cells (clusters c3, c5, c9 and c10; 1,439 cells; **Figures 2B** and **S5A-C**). Interestingly, a much smaller immune cluster (c7, 360 cells), predominantly limited to the DIV6 dataset (**Figures S5B**), in contrast to the zebra finch which had a much higher immune cell proportion (**Figure S2C**). These cluster cell types were concordant to those found in an *in vivo* chicken HH28 gonadal dataset^29^, and the four germ cell clusters all mapped back equally to the PGC chicken germ cells (**Figure S3B**). Compared to the somatic clusters, the collective PGC clusters were positively marked by 8,490 unique DEGs (**Table S6**; a subset is shown in **Figure S5D**). Between the germ cell clusters, only 1,486 DEGs were identified (**Table S7**); many of these DEGs corresponded to their respective inferred sex (c3 and c9 vs. c5 and c10; **Figures S5C** and **S5E**) or predominant cell cycle phase (c3 and c5 vs. c9 and c10; **Figure S5C**).

**Figure 2.**
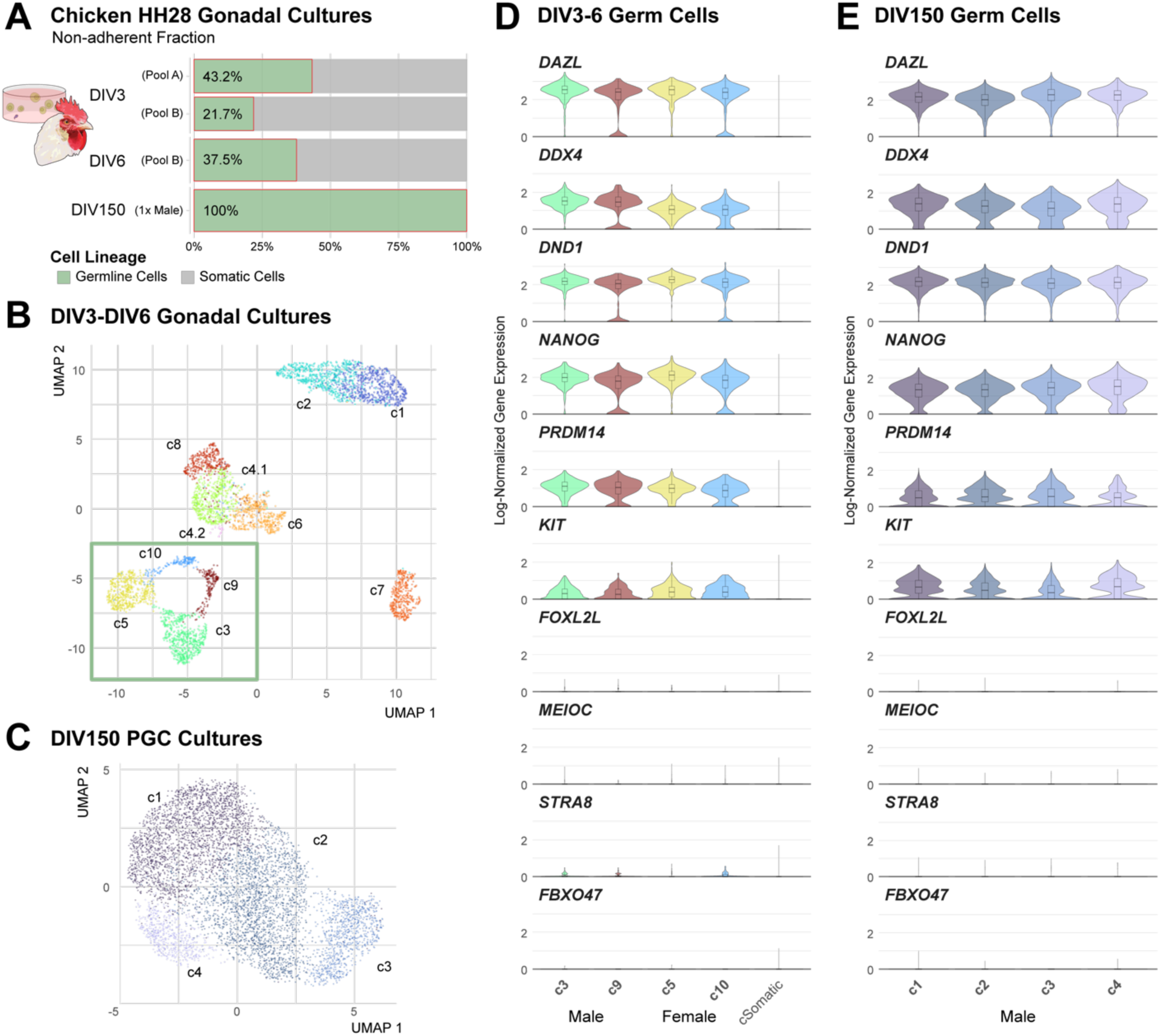
scRNAseq profiling of non-adherent cell types from short- and long-term chicken gonadal cultures. (**A**) Relative proportion of cell types determined from snRNA-Seq analysis among the non-adherent cells in DIV3, DIV6 and DIV150 chicken gonadal cultures. (**B**) UMAP dimensional reduction of DIV3-6 non-adherent chicken gonadal cell cultures, colored and labeled by nearest-neighbor assigned cluster. The green box highlights germ cell clusters. (**C**) UMAP dimensional reduction of DIV150 non-adherent chicken gonadal cell cultures, colored and labeled by nearest-neighbor assigned cluster. Note that all cells in this dataset are transcriptomically assigned as germ cells. (**D**) Select gene marker expression split by cluster for DIV3-6; and (**E**) DIV150 non-adherent cultured cells demonstrating robust early PGC marker expression. Note the consistent expression of PGC markers across all germ cell clusters, and not markers of germ cell differentiation.

In contrast, marker-based cell type inference showed that the DIV150 dataset was entirely composed of germ cells, including those removed by quality filtering (**Figures S6A-B**). After quality filtering, 6,141 PGCs remained (**Table S2**) and resolved into four nearest-neighbor clusters (**Figure 2C**). Between these clusters, 989 DEGs were variably expressed (**Figure S6C** and **Table S8**). Several DEGs were related to germ cell organization and maintenance (*DMRT1*), cell signaling activity (*FGFBP3*, *HEY2*, *RSPO3*), oxidative stress (*EXFABP*, *IPCEF1*), and cell migration (*CXCR4, ROBO1,* and *MSX1*). However, most DEGs were expressed at some level in each cluster and varied in expression level, suggesting that DIV150 PGCs represented a population cycling between homeostatic states rather than undergoing irreversible cell type differentiation.

Importantly, canonical PGC markers^37–40^ were robustly expressed in both DIV3-6 short-term and DIV150 germ cell clusters, including *NANOG, PRDM14*, and *KIT* (**Figures 2D-E** and **S5F**). In contrast, DIV3-6 germ cells expressed low-level PGC differentiation markers like *STRA8* and *FOXL2L* (**Figures 2D** and **S5F**), and no notable expression was detected in the DIV150 culture (**Figure 2E**), in stark contrast to higher levels of these genes in zGC2 and zGC3 DIV6 cells (**Figures 1C**-**G**). This data aligns with another scRNAseq study of chicken PGC cultures^35^, which identified proliferative and homeostatic pathway differences as the primary source of heterogeneity in chicken PGC cultures, rather than germline differentiation. Overall, our findings highlight the maintenance of PGC identity in chicken gonadal germ cells using high-serum culture conditions, unlike in the zebra finch.

#### 3.2.2 Adherent chicken gonadal culture fraction

The chicken DIV3 adherent cells scRNAseq dataset was generated using the same wells as the DIV3 non-adherent sample from Pool B. We did not process the adherent MEF feeder layer from the DIV150 culture, as mitomycin C treatment inhibits transcription^41^. Quality filtering of the adherent DIV3 dataset yielded 1,998 cells across six clusters (**Figure S7A** and **Table S2**). As in the zebra finch adherent cells, four of the six chicken adherent clusters predominantly expressed interstitial cell type markers (**Figure S7B-D** and **Table S9**), along with some IM Progenitor cell types. Small immune (c5; 69 cells) and germ cell (c6; 48 cells) clusters were also identified. Reference mapping of the chicken adherent cells to chicken HH28 gonadal scRNAseq datasets^29^ similarly validated these cell type identities (**Figure S7E**). The chicken germ cell cluster (c6) demonstrated robust expression of both *DAZL* and *NANOG*, the predominant germ cell markers in the non-adherent datasets, with little to no expression of *FOXL2L* (**Figures S7D** and **S7F**). Low-level *STRA8* expression was identified in a small percentage of cells (**Figure S7D**), though was co-expressed with *NANOG* (**Figure S7F**) and did not suggest distinct, differentiating germ cell populations in the adherent fraction.

### 3.3 PGC marker expression persists in zebra finch gonadal and blood cultures

#### 3.3.1 Bulk RNAseq time course of zebra finch gonadal cultures

While the scRNAseq datasets identified a proportional decline of PGCs in the zebra finch non-adherent cell cultures, we wondered whether PGC populations would deplete entirely or persist in low numbers as other populations increased. Accordingly, we generated a bulk RNAseq time course of DIV1-10 zebra finch gonadal cultures to resolve PGC marker expression at the population level. Twelve samples were collected from the non-adherent layer each day (**Table S10**), with additional replicates generated for DIV9 and DIV10. DIV7 and DIV8 were excluded from final analysis, as their samples had low library yields and demonstrated expression profiles that were identified as outliers in PCA and Euclidian distance cluster analyses (**Figure S8**). With the remaining ten samples, Euclidian distance calculations and PCA validated a trajectory along the time course (**Figure 3A-B**).

**Figure 3.**
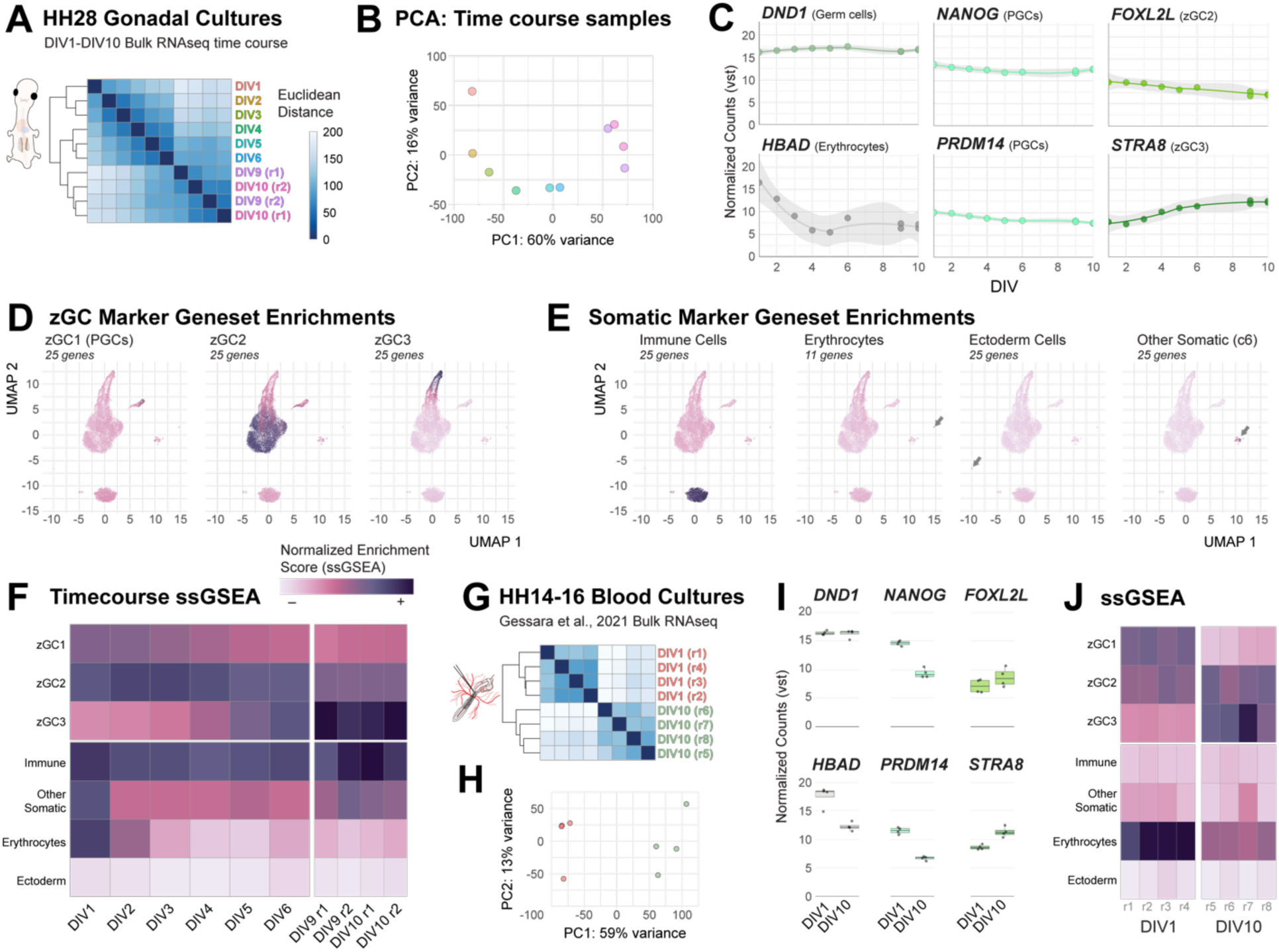
Bulk RNAseq profiling of zebra finch embryonic gonadal and blood cultures over time. (**A**) Heatmap of Euclidian sample distance measurements and (**B**) PCA plot for vst-normalized read counts in bulk RNAseq datasets of a zebra finch gonadal culture DIV1-DIV10 time course. (**C**) Select gene markers of cell types in gonadal culture datasets. (**D**) Heatmap of ssGSEA scores of germline cell type genesets, overlaid onto a UMAP dimensional reduction of the zebra finch non-adherent scRNAseq dataset. (**E**) Heatmap of ssGSEA scores of somatic cell type genesets, overlaid onto a UMAP dimensional reduction of the zebra finch non-adherent scRNAseq dataset. (**F**) Single sample gene set enrichment analysis (ssGSEA) heatmap of zebra finch non-adherent cell type marker genesets for gonadal cultures. (**G**) Euclidian sample distance heatmap and (**H**) PCA plot for previously generated bulk RNAseq datasets of blood PGC cultures at DIV1 and DIV10. (**I**) Gene expression levels the blood PGC cultures, characterizing the same gene markers as in panel C. (**J**) The same type of analyses as in panel F for blood PGC cultures. Note the decrease, but not total depletion, of zebra finch zGC1 geneset enrichments in both culture conditions.

Assessing individual genes in the gonadal bulk RNAseq datasets (**Table S11**), we noted high and persistent expression of the germ cell marker *DND1* (**Figure 3C**). In contrast, the erythroid red blood cell marker *HBAD* declined rapidly, consistent with cells that have erythrocyte morphology dying in just a few days in our PGC culture conditions. Interestingly, both zGC1 and zGC2 markers slightly declined (e.g., *NANOG, PRDM14,* FOXL2L; **Figure 3C**), but were not entirely abolished. Conversely, zGC3 markers robustly increased (e.g., *STRA8* and *FBXO47*), suggesting gonadal culture conditions favored expansion of the zGC3 type.

To better infer the population structure of cell types in these cultures, we selectively identified gene markers for each cell type in the non-adherent scRNAseq dataset (**Table S12**) and assessed the specificity of these expression profiles for single-sample geneset enrichment analysis (ssGSEA) (**Figures 3D-E**). We then mapped these genesets onto the bulk RNAseq samples (**Figure 3F**), which further supported the gradual decline of zGC1 (PGC) and zGC2 marker expression in favor of zGC3 between DIV1-DIV10. Immune cell marker expression was consistently high throughout the time course, and variability in geneset expression could be seen between the DIV9 and DIV10 replicates, indicating some well-to-well heterogeneity even within pooled samples. Nonetheless, the expression level of the zGC1 geneset stabilized around DIV5 at a consistently higher level than the expected low profiles of ectoderm cells or erythrocytes, suggesting that zebra finch PGC populations are present in gonadal cultures as late as DIV10.

#### 3.3.2 Bulk RNAseq assessment of zebra finch blood PGC cultures

Zebra finch PGCs, cultured from HH14-16 embryonic blood in low serum conditions for up to ten days and lentivirally-transduced, were reported to derive transgenic zebra finch offspring^24^. As blood PGCs are isolated during active migration to the gonad, these cultures are likely seeded with a more robust population of migratory-competent germ cells for enhanced germline transmission after injection into embryonic hosts. To compare between gonad and blood-derived PGC culture transcriptomes over time, we utilized a previously generated bulk RNAseq datasets of blood PGC cultures at DIV1 (blood pooled from 5 embryos; 4 replicates) or DIV10 (blood from one male embryo; 4 replicates)^24,42^. Following read mapping and normalization, we found that the samples distinctly segregated the timepoints by both Euclidean distance measurements and PCA (**Figures 3G-H**).

Surprisingly, we found that individual gene marker expression differences between DIV1 and DIV10 zebra finch blood PGC cultures mirrored those found in the gonadal culture time course (**Figure 3I** and **Table S11**). A more comprehensive mapping of the scRNAseq marker genesets by ssGSEA similarly implied extensive germline differentiation in these cultures (**Figure 3J**), with notable zGC2 geneset expression seen even in the DIV1 samples. Interestingly, these samples demonstrated a more dramatic reduction in PGC marker expression between DIV1 and DIV10 than seen in the gonadal time course but was not entirely abolished. This finding is counterintuitive, as blood PGC cultures have been effectively used germline transmission studies in songbirds^24^.

As expected, the blood PGC samples had much higher erythrocyte expression than the gonadal time course (**Figures 3F** vs **3J**) and was not entirely lost by DIV10. Other somatic cell enrichments were also lower in the blood PGC cultures but varied slightly between replicates. Overall, these data show that both zebra finch gonadal and blood PGC cultures undergo rapid and extensive germline differentiation *in vitro* but still retain small PGC populations by DIV10.

### 3.4 Comparison of gonadal migration efficiency between chicken and zebra finch gonadal PGC cultures

Our transcriptomic findings suggested that zebra finch gonadal cultures retained PGC populations. To compare the migratory efficiencies of the zebra finch gonadal cultures, we prepared non-adherent cells at different DIV time points for injection into the blood system of species-specific host embryos at HH14-16 and then assay migratory presence of donor PGCs in host gonads at HH28-30 (**Figure 4A**). The DIV150 chicken PGC cell line (**Figure S1C**), utilizing established long-term culture conditions, was used as a benchmark for optimal migratory efficiency.

**Figure 4.**
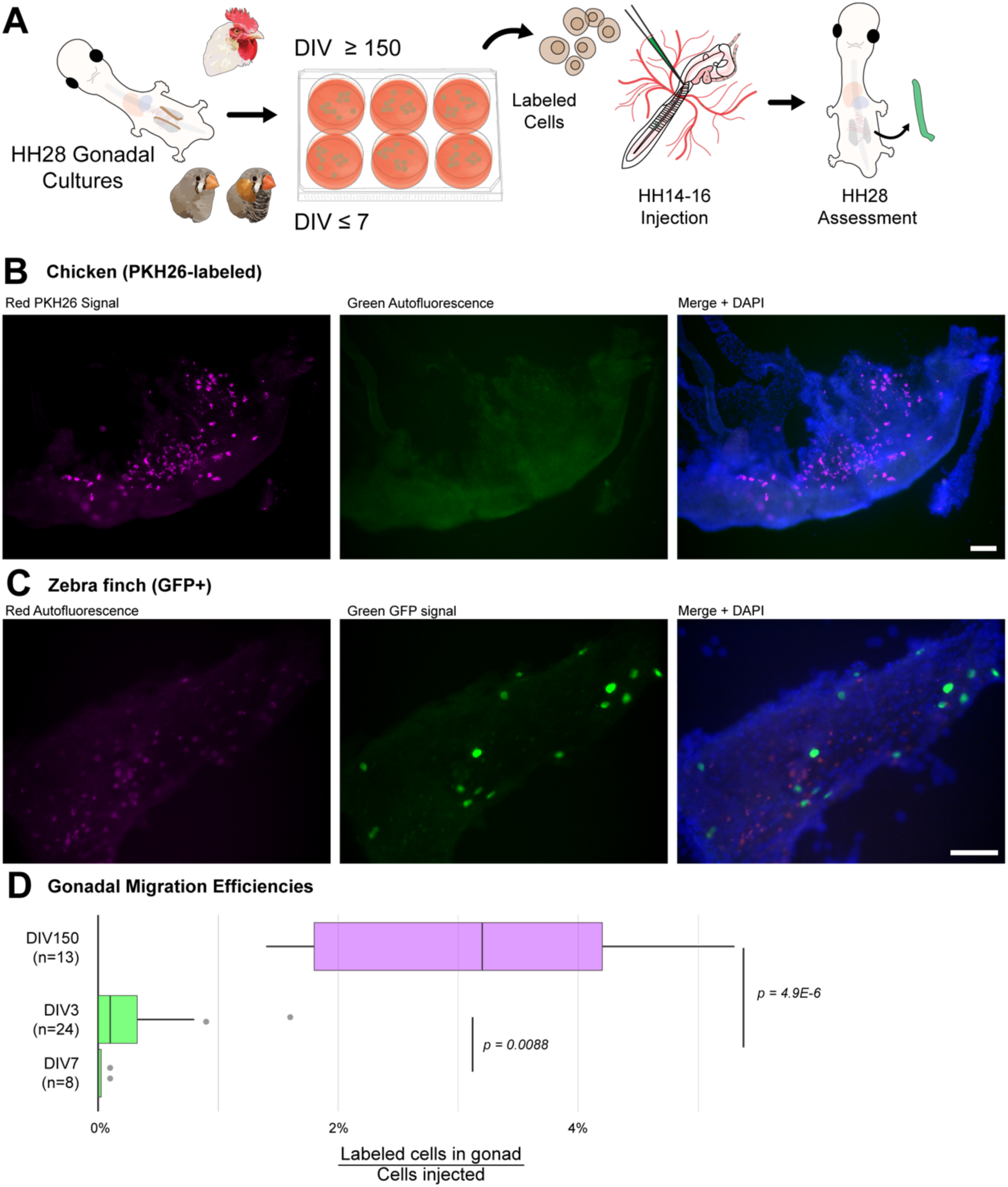
Comparison of migratory efficiency of chicken and zebra finch gonadal PGC cultures. (**A**) Schematic of PGC migration assay. Donor PGCs were cultured for short- (zebra finch) or long-term (chicken) periods, labeled and injected into the blood stream of a host embryo. Host embryonic gonads were then extracted 3 days later to quantify donor PGC migration. (**B**) Exemplar image of dissected ED6.5 chicken gonad containing PKH26-labeled (violet, shifted from red) donor PGCs. Blue labels DAPI nuclear stain. Scale bars = 50µm. (**C**) Exemplar image of dissected ED6.5 zebra finch gonad with GFP+ (green) donor PGCs. (**D**) Host gonad reconstitution efficiency comparison between of PKH26-labeled chicken and GFP+ zebra finch gonadal cultured PGCs, calculated as the number of cells identified in a single gonad out of the estimated number of cells injected at HH14-16.

We labeled cells from the non-adherent fraction of these cultures with PKH26 lipophilic tracking dye prior to injection into HH14-16 host chicken embryos. After a 72-hour incubation (developing to HH28), we found clear and robust PKH26 labeled donor PGCs in the host gonads (**Figure 4B**). No signal in the red channel was identified in control gonads from uninjected chicken embryos (**Figure S9**). However, for zebra finch PGC cultures labeled with PKH26, we noted that with long imaging exposures, zebra finch gonads had autofluorescent cells in injected and uninjected zebra finch embryos (**Figure S10A**), while short imaging exposures showed discontinuous cell staining that made donor cell identification more ambiguous than in the chicken (**Figure S10B**). To mitigate these issues, we cultured PGCs derived from transgenic zebra finches developed by lentiviral integration of hUbC-EGFP+^40^. Compared to the PKH26-labeled zebra finch PGCs, GFP+ PGCs injected into HH14-16 wildtype zebra finch showed a much more consistent and quantifiable signal in the embryonic host gonad (**Figure 4C**).

Across multiple injections using ≥DIV150 chicken (n = 13) or DIV3-10 GFP+ zebra finch gonadal cultures (n = 32), we counted the number of non-autofluorescent, labeled cells per gonad and the percent reconstitution based on the recorded number of PGCs injected into each host (**Table S13**). Injections of long-term chicken PGC cultures consistently resulted in 50-100 donor cells in each of the dissected HH28-30 gonads, corresponding to a 2-5% reconstitution rate (**Figure 4D**). In the zebra finch, the proportion of incorporated cells from short term DIV3-4 cultures was 15-fold less, corresponding to a 0.19% reconstitution rate (**Figure 4D**). By DIV7, zebra finch reconstitution rates were nearly 8-fold lower than DIV3, at 0.03% (**Figure 4D**). Despite these low rates, the level of reconstitution was still non-zero in multiple instances and corresponded to the low-level maintenance of PGC marker expression through DIV10.

### 3.5 Zebra finch gonadal cultures are sufficient for germline transmission

We next sought to assess the viability of short-term gonadal cultures for germline transmission. We chose to pursue a design where we could readily observe germline transmission through feather color, using zebra finches with a recessive chestnut flanked white (CFW) allele as embryonic hosts for donor PGCs cultured from zebra finches with the dominant natural grey (NG) or other non-CFW alleles (**Figure 5A**). A CFW breeding pair will only produce CFW offspring naturally^43^, so any NG offspring would necessarily derive from donor PGCs.

**Figure 5.**
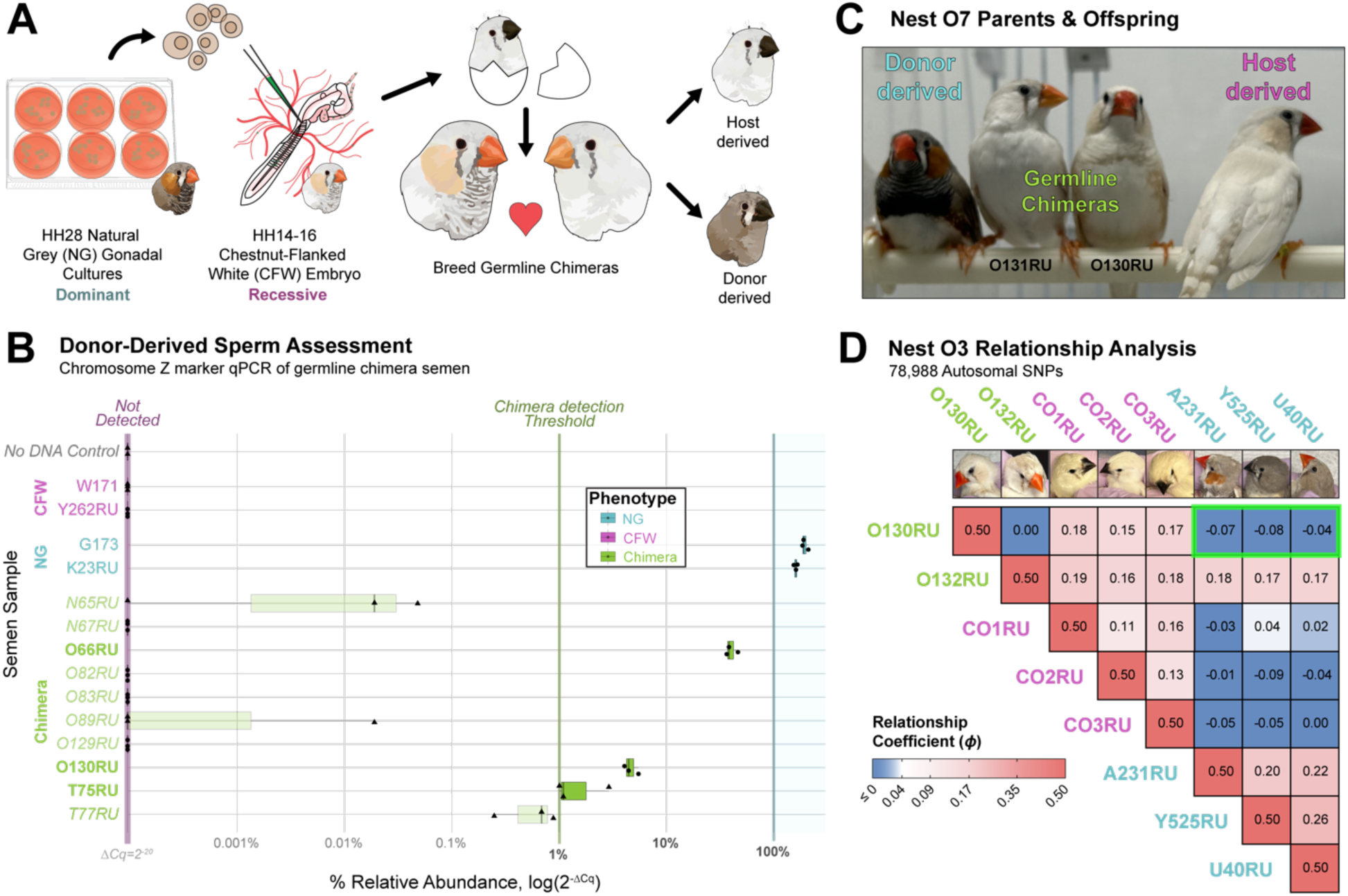
Assessment of germline transmission from zebra finch gonadal PGC cultures. (**A**) Schematic of germline transmission assessment. Short-term zebra finch embryonic cultured PGCs from a natural grey (NG) feathered animal are injected into the embryonic blood stream of an egg from a chestnut-flanked white (CFW) white feathered animal. Germline chimeras are hatched and bred to screen for feather color in offspring. (**B**) Chromosome Z targeted qPCR of zebra finch semen samples, using the NG-marker primer set 68del and the control Chromosome Z primer set 43ctrl. Undetected NG signal was calculated with a ΔCq of 20. A relative abundance threshold of 0.4% (ΔCq ≥ 8) was used to identify donor-sperm signal. Label and boxplot colors represent phenotypic category. Points denote individual well values, with shapes representing different qPCR plates. NTC = No template control. (**C**) Image of Nest O7 parents (Table 1) with exemplar donor (NG)- and host (CFW)-derived offspring. (**D**) Relationships determined by whole genome sequencing analysis for Nest O3 parents and offspring, assessing 78,988 SNVs from Manichaikul et al., 2010. The mother (O132RU) is related to all offspring, while the father (O130RU) is unrelated to the NG offspring (green box).

**Table 1.** Phenotypic assessment of germline chimera offspring.

| Nest | Father | Mother | Hatched Eggs | Feathered Offspring | Host-derived (% CFW) | Donor-derived (% NG) |
| --- | --- | --- | --- | --- | --- | --- |
| O2 | O129RU | O131RU | 62 | 50 | 50 (100%) | 0 (0%) |
| O3 | O130RU* | O132RU | 43 | 38 | 35 (92%) | 3 (8%) |
| O4 | O82RU | W81RU | 38 | 31 | 31 (100%) | 0 (0%) |
| O5 | O83RU | T78RU | 39 | 28 | 28 (100%) | 0 (0%) |
| O6 | T77RU* | T91RU | 38 | 32 | 32 (100%) | 0 (0%) |
| O7 | O130RU* | O131RU | 39 | 24 | 20 (83%) | 4 (17%) |
| O8 | O66RU* | Y549RU† | 0 | 0 | - | - |
| O9 | T75RU* | Y421RU† | 8 | 6 | 6 (100%) | 0 (0%) |
| O10 | N65RU | Y604RU† | 11 | 11 | 10 (100%) | 0 (0%) |
| O11 | O89RU | Y529RU† | 5 | 2 | 2 (100%) | 0 (0%) |
| O12 | N67RU | Y531RU† | 17 | 8 | 8 (100%) | 0 (0%) |

\* NG Sperm Detected
† Uninjected CFW mate

To further validate this experimental design and quantify donor and host gamete proportions in male chimeras, we identified genomic loci associated with the feather color differences between NG and CFW individuals. The CFW allele is known to have a sex-linked inheritance pattern^43^, likely corresponding to a mutation located on the Z chromosome (ChrZ). As part of another study, we performed whole genome-sequencing from 14 CFW and 15 NG individuals (**Table S14**), aligned the raw reads to a high-quality zebra finch reference genome (bTaeGut1.4.pri, GCF_003957565.2), from which we identified two large deletions in ChrZ, one in an intergenic region around 50.83Mb (50del) and the other 68.61Mb (68del) in an intron of *SH3GL2* in all CFW individuals and no NG individuals (**Figure S11A**). A region with equivalent coverage around 43.63Mb (43ctrl) was selected as a control, proximal to the *SLC45A2* gene that is associated with leucistic phenotypes in birds^44–46^. Genomic qPCR on blood samples using primers within the deleted regions only yielded signal from NG animals, and at the similar rate as a control locus present in both phenotypes (**Figure S11B**). The 50del locus produced low level signal in a concentration-dependent fashion, while the 68del locus was not detected at low or high DNA concentrations. To assess the sensitivity of this assay, we serially diluted NG DNA into CFW DNA samples, reliably detecting 1% NG DNA (**Figure S11C**).

With a robust genotype assay developed, we cultured NG gonadal cells for up to 10 days before injecting between 3,000 to 20,000 cells from the non-adherent fraction into the dorsal aorta of CFW host embryos. We hatched 37 putative germline chimeras from 180 injections (20.5%), 17 of which were successfully fostered and matured to breeding age (10 males and 7 females). As expected, each chimera presented a clear CFW phenotype. We assessed DNA samples from sperm of the 10 males and detected NG DNA signal in 6 of them, 3 of which (O66RU, O130RU, T75RU) were above our relative abundance threshold of 1% (**Figure 5B**). O130RU and O66RU had notably high relative abundances (4.6% and 40.3%, respectively), suggesting the robust gonadal integration of donor PGCs following embryonic injections. In somatic DNA from blood or feathers of these same chimeric animals, low-level amplification was detected in a few individuals at a relative abundance ≤1% (ΔCq ≥ 8; **Figure S11D**). However, none of the chimeras with donor-positive ejaculate samples had any detectable 68del amplification in their somatic DNA samples. Male and female chimeras were bred in single-pair cages (**Table 1**), with 7 out of 12 matings producing offspring that survived to feather development (≥7-10 days post-hatch) for feather phenotype screening. Of the three male chimeras with confidently detected donor sperm, only O130RU reliably produced offspring, siring 62 birds screened over two mate pairings. From this CFW host male, his Nests O3 and O7 (**Table 1** and **Figure 5C**) yielded 7 NG offspring (11.3%). No NG offspring have been detected from other chimera nests.

To further validate the parental relationship of NG offspring, we generated whole genome sequencing from blood of the Nest O3 parents (O130RU and O132RU), the 3 NG offspring (Y525, A231, and U40). Three control CFW offspring from Nest O3 were also sequenced. We validated the presence of the CFW-associated deletions (50del and 68del) in the parental CFW chimeras and their CFW offspring (**Figure S11A**). Genome sequences from the NG offspring aligned to the deletion locus. To comprehensively assess genetic similarity between the chimera parents and their offspring, we calculated relationship coefficients (phi)^47^ using 78,988 single-nucleotide variants (SNVs) between Nest O3 individuals and the 29 sequenced individuals as unrelated background controls (**Figure 5D**). For the Nest O3 individuals, we found that the relationship coefficient between the CFW offspring had similar robust relationship coefficients to each parent (phi = 0.13-0.19), while the relationship coefficient for each NG offspring was roughly equivalent to the CFW group for O132RU (phi = 0.14-0.19), but significantly lower to O130RU (phi = -0.1). None of the other comparisons between background samples and the Nest O3 offspring predicted other relationships (*data not shown*). Collectively, these data demonstrate the successful derivation of offspring from short-term zebra finch cultured PGCs.

## 4 Discussion

Our study reveals key transcriptomic and developmental differences that can explain why chicken gonadal PGC cultures are more viable for assisted reproductive techniques than from zebra finch PGC cultures. While the germline proportion of short-term gonadal cultures were much higher in zebra finch compared to chicken, only the zebra finch germ cells rapidly differentiate and dramatically reduce the PGC proportion after a few days. This extensive germline differentiation occurs in both gonadal- and blood-derived PGC culture protocols, despite the latter demonstrating robust germline transmission in a previous study^24^. Despite these critical species differences in germline differentiation and long-term culture viability, we showed that zebra finch gonadal cultures retained a sufficient population of PGCs or other migration-competent germ cells that could colonize the gonads of host embryos, produce sperm in adult chimeras, and subsequently give rise to donor-derived offspring. In demonstrating this potential, we developed additional assays to compare migration competence, genotype gametic proportions, and validate the relationship of offspring using clear phenotypic markers and robust sequencing techniques. Many of these genomic and reproductive methods will be valuable to the avian biology community.

The higher germline proportion in early zebra finch cultures relative to chicken reflects the abundance of germ cells *in vivo* for each species^26^. Additionally, we noted that the chicken cultures had much higher cell death rates in short-term cultures compared to finch, possibly due to the extended trypsin dissociation required for the larger tissue or species-specific differences in shear stress tolerance. Despite these differences, only the chicken cultures have been capable of long-term maintenance and self-renewal in culture. We confirmed in both species that the non-adherent cell fraction contained the majority of PGCs in these cultures, while the adherent cell fraction had few germ cells that did not represent distinct populations. In contrast, the adherent cell culture fractions in each species consisted mostly of somatic cells derived from an interstitial and mesodermal lineage^48^, though cross-species comparisons suggest that these populations are more developmentally mature in the zebra finch gonad by HH28^29,49^. While differentiation appears to occur even in feeder-free blood PGC conditions, one possibility may be that the gonadal somatic cells of chicken better support the survival and maintenance of PGCs prior to self-renewing transformation. Further comparative assessments of adherent population signaling may provide additional insights on the derivation of long-term PGC cultures in other species.

The heterogeneity of the zebra finch PGC cultures also offers a look at the dynamic profile of germline development in songbirds, including genetic contributions from the GRC, a unique and enigmatic genomic element not found in other jawed vertebrates (*Gnathostomata)*^20,34^. As previously characterized *in vivo*^29^, we similarly identified robust *in vitro* expression of identified GRC genes^21,50^, several of which were differentially expressed between all three germline cell types in culture (zGC1-3). This robust GRC gene expression also suggests that each germ cell type retains this chromosome and would be a viable substrate for comprehensive biobanking of songbird genetic material. While the GRC sequence remains largely incomplete, our work highlights the potential value of *in vitro* models in exploring the GRC’s functional role in the development and maintenance of the songbird germline.

More broadly, our short-term cultures offer another comparative model for vertebrate germline development. Derived from the HH28 gonad, our zebra finch cultures contained both the *NANOG+* zGC1 (PGCs) and differentiating *FOXL2L*+ zGC2 populations found *in vivo*^29^. The zGC2 populations also maintained high rates of mitotic marker gene expression *in vitro*, suggesting a highly proliferative state resulting in zGC2 expansion, which may partially explain the proportional decline in PGCs over time. However, we also noted strong enrichments of zGC2-associated geneset expression in blood PGC cultures, suggesting that all current zebra finch PGC culture protocols at least partially enable PGC differentiation into zGC2 populations. In both zebra finch culture methods, a more clustered morphology has been noted compared to chicken^24,26^. These clusters mirror nest morphologies implied by histological characterizations of the HH28 zebra finch gonad^29^, and are suggestive of *FOXL2L*+ germ cell nests characterized in other vertebrate clades^51,52^. Interestingly, the eventual decline of the zGC2 populations in favor of *STRA8+* zGC3 populations in both gonadal and blood PGC datasets unexpectedly identified a germline identity not found *in vivo* at HH28. In many vertebrate lineages, *STRA8* encodes a critical driver for retinoic acid (RA)-mediated meiotic transition^53–55^. In the HH36 female gonad, *STRA8* expression is also detectable in differentiating germ cells, coinciding with the onset of *FOXL2L* expression^29,30^ that precedes pre-meiotic DNA synthesis^31^. In the zebra finch gonadal cultures, we found that *FOXL2L* is most highly expressed in zGC2 and declines as *STRA8* is upregulated in zGC3. Further work will be needed to characterize the developmental trajectories of these decoupled genes *in vivo* and explore the diversity of germline gene networks across the avian clade.

Our experiment showing NG-associated donor DNA in CFW host sperm, reliably detected in three male chimeras, is promising. One of these germline chimeras (O130RU) has generated seven F1 NG offspring from two mate pairings. While relatively high donor DNA signal was detected in two other males, breeding did not produce a significant number of offspring, highlighting one of many difficulties in assessing germline transmission in non-poultry avian species. As of publication, these three germline chimeras are alive and paired with mates to assess germline transmission. In the study by Gessara et al^24^, DIV10 PGC cultures from blood were able to generate 10 germline chimeras that produced donor-derived offspring out of 22 injected eggs. Notably, these chimeras were generated by injecting cultured germ cells into recently laid blastodiscs, rather than into the vasculature of HH14-16 embryos. While our re-analysis of their *in vitro* transcriptomic data identified extensive germline differentiation, the integration of cultured cells at this earlier stage may provide a more suitable environment that promotes pluripotent and migratory pathways in PGCs and potentially in differentiated germ cells. While our study demonstrates that gonadal cultures are a viable alternative for germline transmission after injection into later embryonic stages, future work will be beneficial to explore the optimal stage for donor cell reintroduction, as well as other aspects of assisted reproduction and avian husbandry, such as artificial insemination to enhance breeding rates^56^.

We believe our approach will be useful for avian biobanking, particularly in applications where transgenic manipulations are not needed. The cost reduction in genome sequencing provides a more accessible and rigorous means to screen for donor-derived offspring, even in the absence of a phenotypic marker. Our methods also provide greater flexibility when collection from narrow embryonic stages is not possible, such as in endangered species conservation. Recent work in chicken has demonstrated that biobanked gonads from a wide range of embryonic stages can be used to derive long-term PGC lines to derive offspring^13,14^. As more than 12% of birds are threatened with extinction^57^, including more than 1,000 passerine species, such approaches beyond the chicken will be critical tools to catalog, preserve, and potentially restore genetic diversity in vulnerable avian populations. Accordingly, we hope that this study will serve as a roadmap for future work in non-model and endangered avian species where developmental staging and gene manipulation resources are absent.

On the other hand, the use of fluorescent markers was valuable to confidently assess the migration efficiencies of zebra finch gonadal cultures. While robust rates of chicken PGC migration could be quantified using lipid membrane dyes, in the zebra finch this approach did not yield consistent signal of individual cells and was confounded by low rates of reconstitution and gonadal autofluorescence. Instead, our use of a lentiviral GFP transgenic zebra finch line^58^ provided a clearer and more uniform signal to detect donor cell integration. This transgenic line has been used in other grafting experiments^59^, and our work further highlights the value of transgenic songbirds for tool development and insights into biological processes.

Overall, our study outlines the scientific and reproductive applications of short-term zebra finch gonadal cultures, while further highlighting areas to enhance efficiency. One area for improvement will be to mitigate PGC differentiation and promote self-renewal *in vitro*, potentially starting from fully defined media conditions as optimized for chicken blood PGC cultures^6,35^. Specifically, the exclusion and replacement of serum in media conditions offers a more comprehensive look at how the survival and self-renewal of PGCs *in vitro* is affected by specific growth factors and other reagent modifications. Another enhancement may be to isolate PGCs from heterogeneous cultures by FACS, providing purer migration-competent populations for injection. This method will benefit from further analyses of scRNAseq datasets and histological validation to identify potential through antibody- or lectin-based targeting approaches^60,61^.

## 5 Methods

### 5.1 Animal husbandry and sources

Zebra finches were maintained under a 12:12-h light/dark cycle at 18-27°C and breeding pairs provided with a finch seed blend, millet spray, egg mash with fresh squeezed oranges daily, and fresh fruits and vegetables once to twice weekly. A hanging nest box and *ad libitum* jute/cotton mix for nesting material were placed in each cage. Eggs were collected daily and stored at 16-18°C, 80% humidity for up to 7 days. Fertile White Leghorn chicken eggs were obtained from Charles River Laboratories. Eggs were incubated in egg incubators (Showa Furanki, Cat. #P-008B) at 37.5-38.0°C, 40-60% humidity) until reaching HH14 (60-72 hours) or HH28 (5.5-6.5 days).

### 5.2 Cell Culture

Gonadal cultures from HH28 zebra finch and chicken embryos were generated as previously described^9,26^, incubated at 37°C and 5% CO_2_ in a Heracell VIOS 160i incubator (Thermo Scientific). Cell culture media recipes used in this study are available in **Table S15**.

#### 5.2.1 Short-term chicken and zebra finch gonadal cultures

For each primary culture pool, forty embryos (unsexed male and female) were pooled and dissociated at 37°C with 0.05% trypsin (zebra finch: 5min; chicken: 15min) and equally split across multiple wells of a 12-well plate (zebra finch = 8 wells; chicken = 12 wells). For the non-adherent fraction, the supernatant was collected from all wells after briefly swirling the plate and pelleted at 300xg for 5min. For the adherent cells, wells were washed twice with HBSS and dissociated with 0.05% trypsin for 5min, neutralized in PGC media, and pelleted at 300xg for 5min. For each fraction, half of the pellets were taken for DIV3 samples, while the remaining cells were resuspended in fresh PGC media and incubated in 4-6 wells until DIV6.

#### 5.2.2 Long-term chicken PGC cultures

To generate a long-term chicken PGC line, gonads from single embryos were cultured as in the short-term gonadal cultures for more than one month before transfer to wells with a feeder layer of CF1 MitC-treated MEFs (Gibco, Cat. #A34959) in the same chicken PGC media conditions. One male chicken PGC line, designated IP6, proliferated for multiple months. At DIV150, we purified the cultures through dead-cell removal by MACS (Miltenyi Biotec, Cat. #130-090-101) and stored cryopreserved cells in a 5:4:1 mix of chicken PGC culture media (**Table S15**), FBS, and DMSO.

#### 5.2.3 Sample Collection

Non-adherent cells were pelleted at 250xg for 3min and resuspended in 0.05% trypsin-EDTA at 37°C (zebra finch: 10min; chicken: 3.5min). Trypsin was inactivated with an equal volume of PGC media before pelleting and resuspension in Sorting Buffer (**Table S15**). Following non-adherent cell aspiration, the adherent cell layer was dissociated with Accutase (Sigma, Cat. #A6964) for 10min. Cells were dislodged by gentle tapping, pelleted at 250xg for 3min, and resuspended in Sorting Buffer. All cell suspensions were filtered through a 40µm cell strainer (Bel-Art, Cat. #03-421-228) immediately before sorting on a BD FACSAria™ II at The Rockefeller University Flow Cytometry Resource Center (RRID:SCR_017694) for dead cell and debris exclusion using DAPI.

### 5.3 Single-cell RNA sequencing

#### 5.3.1 Single Cell Capture on 10x Genomics Chromium

Sorted samples were immediately processed for scRNAseq, as previously reported^29^. For the DIV150 chicken cells, a cryovial was thawed and cultured for at least one week prior to collecting the cell suspension. Appropriate volumes of water and cell suspension to yield approximately 7,000 cells per sample, following the manufacturer’s instructions for Chromium Single Cell 3ʹ GEM, Library & Gel Bead Kit (v3). The droplet emulsion was reverse transcribed at 53°C for 45 min before heat deactivation at 85°C for 5 min and cDNA amplification. cDNA was measured on a Qubit Fluorometer (ThermoFisher #Q33238) and validated for size and purity on an Agilent Fragment Analyzer (#M5310AA) using the High Sensitivity NGS Kit (Agilent, Cat. #DNF-474-0500).

cDNA samples were normalized (zebra finch: 100ng/sample; chicken: 50ng/sample) for Illumina sequencing library preparation following the 10x Genomics protocol (Chromium Single Cell 3’ Reagent Kit v3). Illumina libraries were quantified using a Qubit Fluorometer (Thermo, Cat. #Q33238) and visualized using an Agilent Fragment Analyzer (#M5310AA, DNF-474-0500). Libraries were labeled using the Chromium i7 Multiplex Kit (10X Genomics, Cat. #PN-120262) and sequenced on a NovaSeq S4 (pair-ended with read lengths of 150bp).

#### 5.3.2 Downstream Analysis

Quality control analysis was performed using similar approaches in our previous study^29^. Briefly, raw sequencing data, was aligned to either the zebra finch or chicken reference genome (depending on the sample) using CellRanger (version 6.0.1). The DIV150 chicken dataset was also aligned to the mouse genome, which showed no sample cross-contamination by the MA-MEF feeder layer in any chicken PGC barcodes (*data not shown)*. Subsequent processing steps were performed using R (version 4.4.1).

Raw and filtered count matrices were run through soupX (version 1.5.2) for ambient RNA assessment and correction. These corrected counts were loaded into Seurat (version 5.1.0) for subsequent processing. As opposed to a universal cutoff for cell barcode trimming, inferred cell-type thresholds were determined for read, feature, and mitochondrial gene values to account for differences hypertranscriptionally activated populations^62,63^. Doublets were identified and removed using DoubletFinder (version 2.0.4).

Filtered samples were merged and integrated to produce the number of principle components for dimensional reduction and clustering resolution (Table S1). In some cases, (e.g., cluster c5 in the zebra finch non-adherent layer), small populations with distinct cell type marker identities were manually split (c5.1: *NANOG+* PGCs; 391 cells vs. c5.2: *SOX10+* ectoderm cells; 11 cells) using the CellSelector function and cross-validated using the FindSubClusters function. Cell sex was inferred using UCell module scores of Chr.Z and Chr.W genes. Barcodes were only assigned male when no ChrW gene expression was detected, while female barcodes were assigned above a certain expression level. We noted that zebra finch germline populations generally had lower expression levels of ChrW genes than somatic populations, and defined thresholds for each group. This difference was not seen in chicken, requiring only one threshold. illustration of this process is found in **Figures S2E** and **S5E**.

DEGs were determined using the FindAllMarkers function in Seurat and filtered using the following criteria: Log_2_ fold-change ≥ 0.5; cluster expression ≥ 10%; Adjusted p-value ≤ 0.05. Tables and figures were generated using standard workflows. A detailed workflow is available through Github (see Data Availability statement).

### 5.4 Bulk RNA sequencing

For the non-adherent time course samples, zebra finch gonadal cultures were collected in a similar manner to the scRNAseq dataset, and RNA was extracted using RNA micro kit (QIAGEN, Cat. # 74034), reverse transcribed using SuperScript Reverse Transcriptase II (Invitrogen, Cat. # 18064014), and PCR amplified for 16 cycles with KAPA Hi-Fi HotStart ReadyMix (KAPA Biosystems, Cat. #KK2601). Sequencing libraries were prepared using the NEBNext® Ultra™ II FS DNA Library Prep Kit for Illumina (NEB, Cat. #E7805S) before being sent to Novogene Corporation, Inc. (Sacramento, CA, USA) for sequencing over two lanes of an Illumina HiSeq 4000. For analysis, gonadal and blood culture samples (**Table S10**) were aligned to the bTaegut1.4.pri reference genome using HiSAT2 (version 2.2.1) and analyzed with DESeq2 using a standard pipeline. Markdown files of the analysis may be found on Github.

### 5.5 Injection of cultured PGCs

#### 5.5.1 HH14 dorsal aorta injection

For PGC culture injections into host embryos, the DIV150 chicken PGC line, thawed from a cryovial and maintained in culture for at least one week. For the zebra finch, wildtype or UbC-GFP+ gonadal cultures were maintained 3-10 days *in vitro* prior to injection. The non-adherent cells were collected and stained with PKH26 dye using manufacturer protocols (Sigma-Aldrich, Cat. #PKH26GL), after which the cells were resuspended in HBSS (Gibco, Cat. #4170112) containing approximately 50mM of Fast Green FCF (Sigma-Aldrich, Cat. #F7252) to validate proper targeting of the vasculature. For the migration assay of the GFP+ zebra finch gonadal culture and chimera generation, gonadal culture supernatants were pelleted and directly resuspended in HBSS with Fast Green FCF. For the generation of different feather color chimeras, wildtype zebra finch cultures from NG nests between DIV4 and DIV10 were pooled, centrifuged at 300xg for 5min, resuspended, and injected 0.3-2µL into the blood of embryo hosts from CFW nests. Depending on yield, the final cell concentration varied between 800-20,000 cells/µL.

For injection surgeries, chicken and zebra finch host eggs were incubated for 2.5-3 days and candled to identify embryos between HH13-16. In chicken, approximately 1mL of albumen was removed and the shell opened through the air sac with curved scissors (Fine Science Tools, Cat. #91461-11) to reveal the embryo. In the zebra finch, a small opening was made directly over the embryonic trunk with forceps, taking care to avoid piercing the shell membrane. The shell membrane was then covered with a small amount of sterile saline and opened to reveal the dorsal aorta. In both species, between 0.3-2µL of cell suspension was injected using a Nanoject III (Drummond Scientific, Cat. #3-000-207) mounted to a stereotaxic micromanipulator (Kopf Instruments, Cat. #961). Following injection, the egg was rotated to attempt to move the embryo away from the opening before sealing with tape (3M, Cat. #600) or silicone mold rubber (Smooth-On Body Double^TM^ Fast Set). Eggs were then artificially incubated for three days and either moved to a foster nest for chimera hatching or opened to assess host gonad reconstitution of cultured PGCs. The cell suspension concentration and injection volume were documented for each injected embryo.

To assess cultured PGC migration, HH28-30 gonads were carefully dissected in a single piece and placed directly into ProLong Gold mounting medium with DAPI (Invitrogen, Cat. #P36931) on a SuperFrost Plus slides (Fisherbrand, Cat. #1255015) and cover slipped (Corning, Cat. #2975224). Gonads were imaged within one hour of dissection to mitigate tissue distortion and autofluorescence, using a Leica DMIL LED inverted microscope with a DFC3000G camera. In some cases, only one gonad was successfully retrieved. To calculate migratory efficiency for each injected embryo, the number of labeled cells in one gonad was divided by the estimated number of cells injected. In cases where both gonads were collected, the gonad with more labeling was quantified. Statistical analysis was performed using the ggpubr package (version 0.6.0) in R.

### 5.6 Assessment of Germline Transmission

#### 5.6.1 Somatic DNA collection and sequencing

DNA was collected from 29 phenotypically catalogued zebra finches as well as the parents and 6 offspring from Nest O3. 50-100µL of blood was collected via the jugular vein (for whole genome sequencing) or 4-5 feather cuticles (for chimeric DNA analysis) and processed using the DNEasy Blood and Tissue Kit (QIAGEN, Cat: #69504). Details on sample identity, library preparation, and sequencing are provided in **Table S14**.

#### 5.6.2 Genomic Analysis

For identification of NG and CFW-associated loci on the Z chromosome, fastq files with raw reads were aligned to bTaeGut1.4.pri with bowtie2^64^ (v2.4.4) and reads were deduplicated with UMIs using Picard^65^ (v2.27.5). Coverage depth across 250bp bins was calculated using mosdepth (version 0.3.3). Regions without any sequence coverage in the CFW cohort and in all NG individuals were visualized in IGV (version 2.17.4)^66^. For relationship analysis, A VCF file was generated using freebayes (v1.3.4) from zebra finch autosomal sequence alignments (**Table S14**), including samples of the Nest O3 parents and offspring. Strict quality filtering was applied using vcftools^67^ (v0.1.16) with the following parameters: read depth ≥ 3, only biallelic variants, Q-score ≥ 30. The resulting vcf file was subsequently used to derive relationship coefficients (φ) using an implementation of <u>k</u>inship-based <u>in</u>ference for <u>g</u>enome-wide association studies (KING) through the vcftools package. Following KING inference guidelines^47^, related individuals (2^nd^ degree relationship or greater) were inferred from pairs with φ ≥ 0.09, while unrelated individuals were inferred when φ ≤ 0.04.

#### 5.6.3 Sperm DNA Preparation

Sperm DNA samples were generated as described in a prior study.^68^ Briefly, semen samples were collected by cloacal massage, placed into 20-50µL PBS, and stored at -20°C. Samples were later quickly thawed and DNA extraction performed using the DNEasy Micro DNA Kit (QIAGEN, Cat. 69504) and eluted in 30µL diH_2_O. DNA concentration was quantified using a dsDNA HS Assay Kit (Invitrogen, Cat. Q32851) on a Qubit3 Fluorometer (Life Technologies, Cat. Q33216). Sample concentrations ranged from 7.5-240ng.

#### 5.6.4 Genomic DNA qPCR

qPCR primers were designed against regions of the genome that difference, using primer-BLAST^69^. Reference (43ctrl) and target (50del or 68del) primer sets were run in triplicate for each sample assessed. In a MicroAmp 96-well reaction plate (Applied Biosystems), 2-40ng of sample DNA was mixed with 5µL 2x PowerUp SYBR Green Master Mix (Applied Biosystems, Cat. A25742), forward and reverse primers to a final concentration of 200nM, and diH_2_O for a total reaction volume of 10µL. Plates were sealed, vortexed, and centrifuged for 2min at 2000xg before running on a StepOnePlus Real-Time PCR System (Applied Biosystems; RRID: RRID:SCR_025855) for 40 cycles (T_m_=60°C). Primer sequences are in **Table S16**. For simulated DNA assessment, blood DNA samples from either an NG or CFW male was diluted to 1ng/µL and validated before 1:10 serial dilution of the NG sample into CFW sample.

Genomic qPCR results were analyzed in R using the tidyqpcr package^70^ (v1.1). A “high-confidence” detection threshold was determined by Cq≤35, ΔCq≤8, and a standard deviation ≤ 0.25 between technical replicates (**Figure S11B-D**).

## 6 Acknowledgments

Thanks to the Rockefeller University Flow Cytometry (RRID:SCR_017694) and Bioinformatics Resource Centers. The authors would also like to acknowledge Gist Croft, Ben Novak, Kyung Min Jung, Jae Yong Han, Carlos Lois, Tarciso Velho, Claudio Mello, and John Bracht for helpful discussions and suggestions in experimental design. Several zebra finch samples for DNA were generously provided by Carlos Lois and Nicole Schweers. Funding for this work was provided by Revive and Restore, the National Science Foundation (EDGE Grant ID 1645199)), The Robert W. Wilson Trust, and the Howard Hughes Medical Institute.

## 7 Author Contributions

M.B., A.K., C.S., A.S., and E.J., conceived the study. T.C., A.K., and E.J. provided instruments, materials, and reagents. M.B., C.S., A.S., E.H., K.B., and A.K., performed the experiments. M.B., E.H., R.M., P.C., and B.H. generated sequencing data. M.B., J-D.L., W.W., and T.C. analyzed the transcriptomic data. M.B., and A.K. analyzed the gonadal migration data. M.B. analyzed the qPCR and genome sequencing data. G.D. and L.T. provided husbandry support and offspring identification. M.B., A.K., and E.J. wrote the manuscript. All coauthors contributed to manuscript revision.

## 8 Data and Code Availability

Accession numbers for all datasets used will be generated after uploading to NCBI and provided in the final manuscript. Previously submitted accession numbers may be found in **Tables S1** (scRNAseq; BioProject TBD), **S10** (Bulk RNAseq; BioProjects PRJNA637305 and TBD) and **S14** (WGS sequencing; BioProject TBD). Final code and markdown file workflows used for bioinformatic analyses will be uploaded to Github (https://www.github.com/Neurogenetics-Jarvis). All other data is available through reasonable request to the corresponding authors.

## 9 Declaration of generative AI and AI-assisted technologies in the writing process

During the preparation of this work the authors used ChatGPT to assist with structuring initial text outlines for clarity. After using this tool, the authors reviewed and heavily edited all texts and take full responsibility for the content of the publication.

## 11 Supplemental Figures

**Figure S1.**
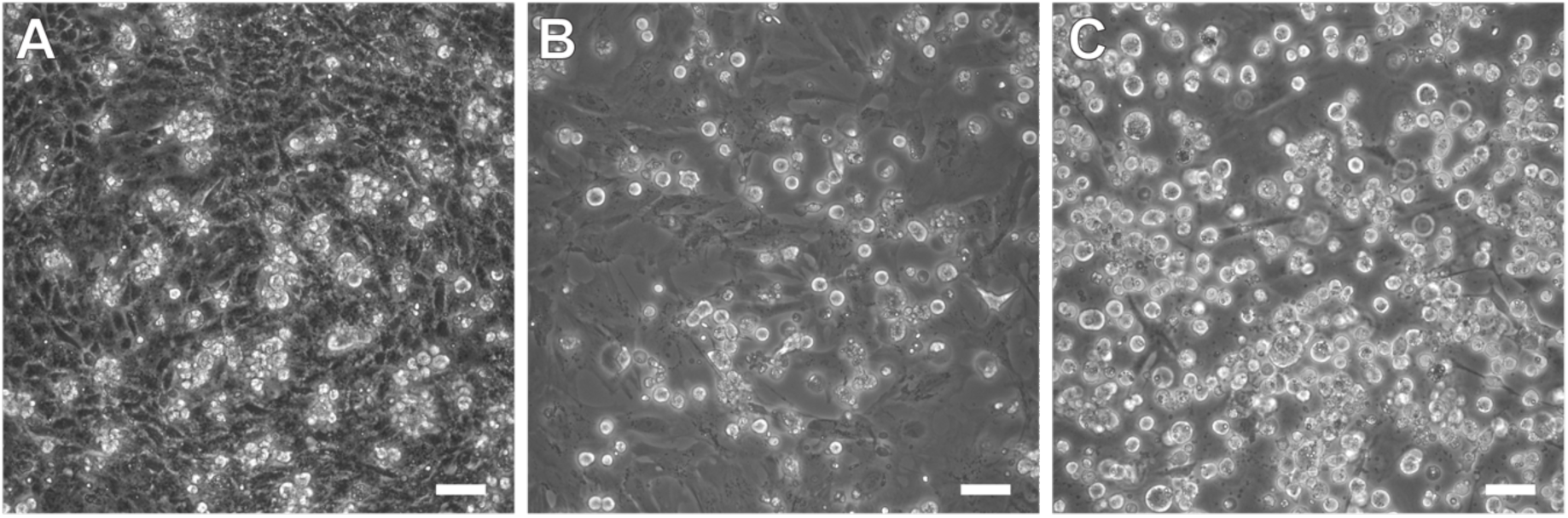
Example images of gonadal cell cultures. (**A**) Short-term (DIV6) zebra finch gonadal culture. (**B**) Short-term (DIV3) chicken gonadal culture. (**C**) Long-term (DIV150) chicken PGC culture line. The chicken culture example shown was thawed from a cryovial and cultured for at least one week on a mitotically-inactivated MEF feeder layer. In all cultures, note that PGCs have characteristic maintain rounded and bright morphologies in the non-adherent fraction (green arrows), while somatic cells are darker and adhere to the plate surface (orange arrows). Note the generally more clustered morphology of the zebra finch non-adherent cells compared to chicken.

**Figure S2.**
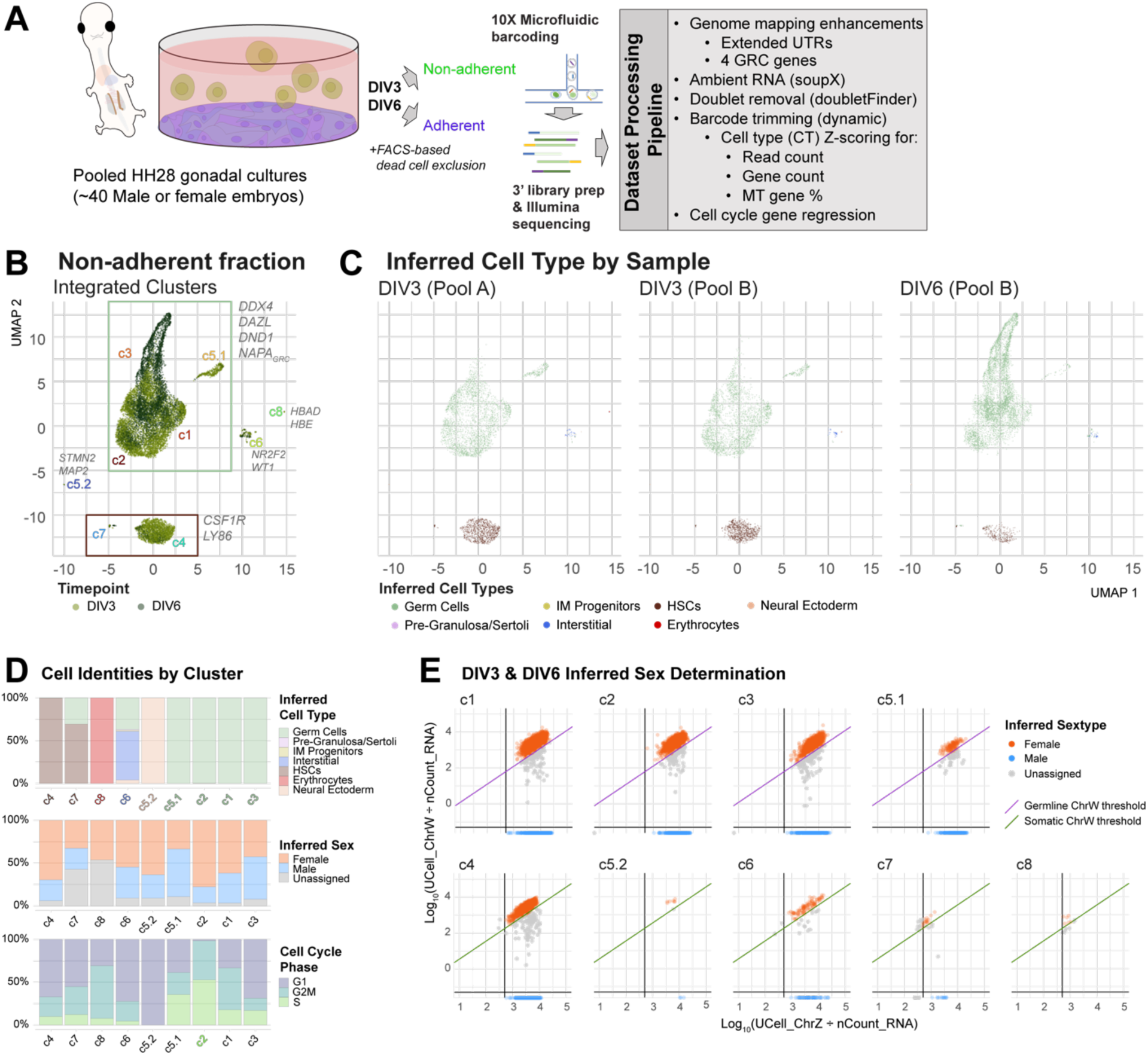
Processing and additional analysis for scRNAseq datasets of zebra finch short-term gonadal cell cultures. (**A**) Schematic of workflow. Embryonic gonads at HH28 are pooled from multiple animals, grown in culture for 3 or 6 days, the non-adherent and adherent cells separated, and scRNAseq performed on each sample using the listed quality control considerations. (**B**) UMAP dimensional reduction of DIV3 and DIV6 integrated non-adherent zebra finch gonadal samples, colored by DIV and labeled by nearest-neighbor cluster assignment. Example cell type marker genes enriched in each cluster are indicated. (**C**) Same UMAP profile colored by inferred cell types at DIV3 and DIV6, and the different pools. (**D**) Proportional bar charts of inferred cell type, sex, or cell cycle phase by cluster. (**E**) Scatterplots of log-normalized Chr.Z (x-axis) and Chr.W (y-axis) geneset expression, split by cluster. Dots denote geneset enrichments for each cell barcode, colored by inferred sex; lines denote inference thresholds for sex determination per barcode.

**Figure S3.**
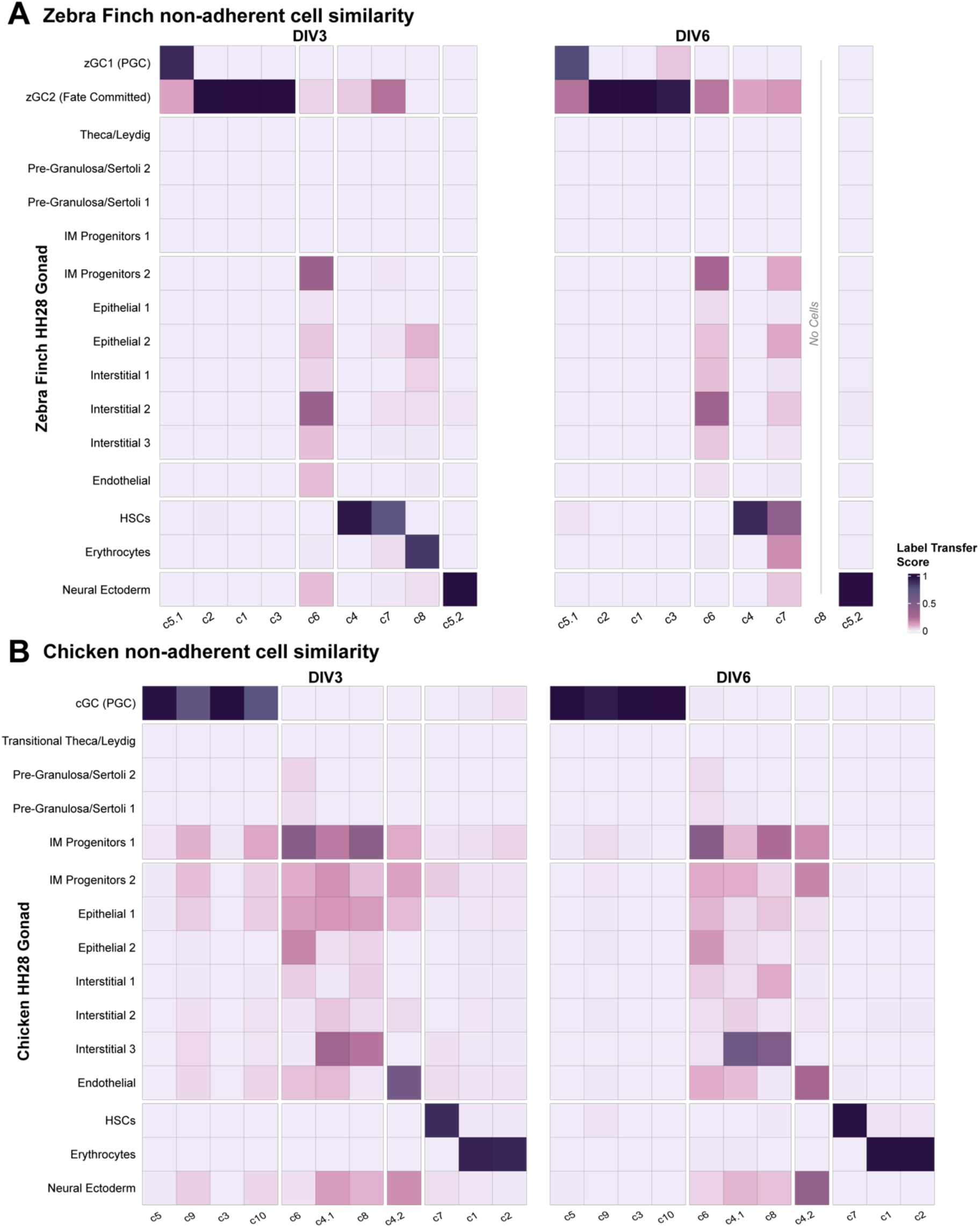
Cell type scRNA-Seq transcriptome comparisons between short-term cultured embryonic gonadal cells and *in vivo* gonadal cells. (**A**) Confusion matrices of Seurat label transfer scores for similarities between zebra finch non-adherent cell types in DIV3 and DIV6 cultures from zebra finch in this study relative to HH28 in-vivo gonadal datasets from our previous study^29^. (**B**) The same analysis applied to the chicken data sets from the current and previous study^29^ at the same time points. Greater similarity is denoted by darker colors.

**Figure S4.**
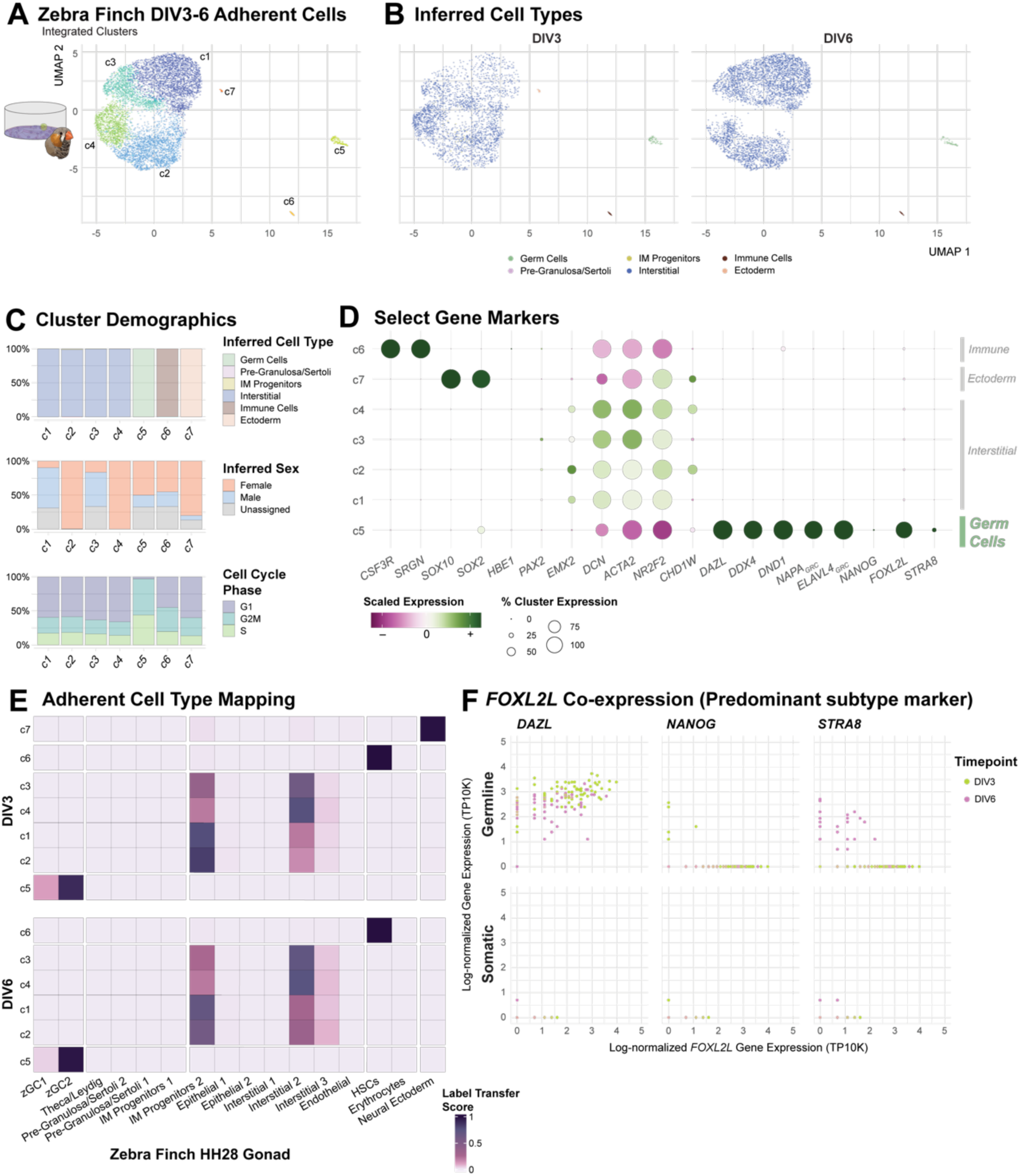
scRNAseq profiling of adherent cell types from short-term zebra finch gonadal cultures. (**A**) UMAP dimensional reduction of integrated adherent zebra finch gonadal samples, colored and labeled by nearest-neighbor cluster assignment. (**B**) UMAP dimensional reductions of each time point, colored by inferred cell type. (**C**) Proportional bar charts of inferred cell type, sex, or cell cycle phase by cluster. (**D**) Dotplot of select gene markers expressed in each cluster, highlighting cell types. (**E**) Confusion matrix of cell type similarities between the zebra finch adherent cultured cells and the zebra finch *in vivo* HH28 gonadal dataset. (**F**) Co-expression analysis of germ cells (cluster c5 from panel A) and somatic cells (all other clusters) from the adherent cells, colored by DIV. The dominant non-adherent germ cell type marker, *FOXL2L,* is overlaid with other markers of zebra finch germ cell types. Note the high *FOXL2L* co-expression with *DAZL* (germ cell marker), but much less than with *NANOG* (zGC1 marker) and *STRA8* (zGC3 marker), suggesting similar germ cell populations as in the non-adherent dataset.

**Figure S5.**
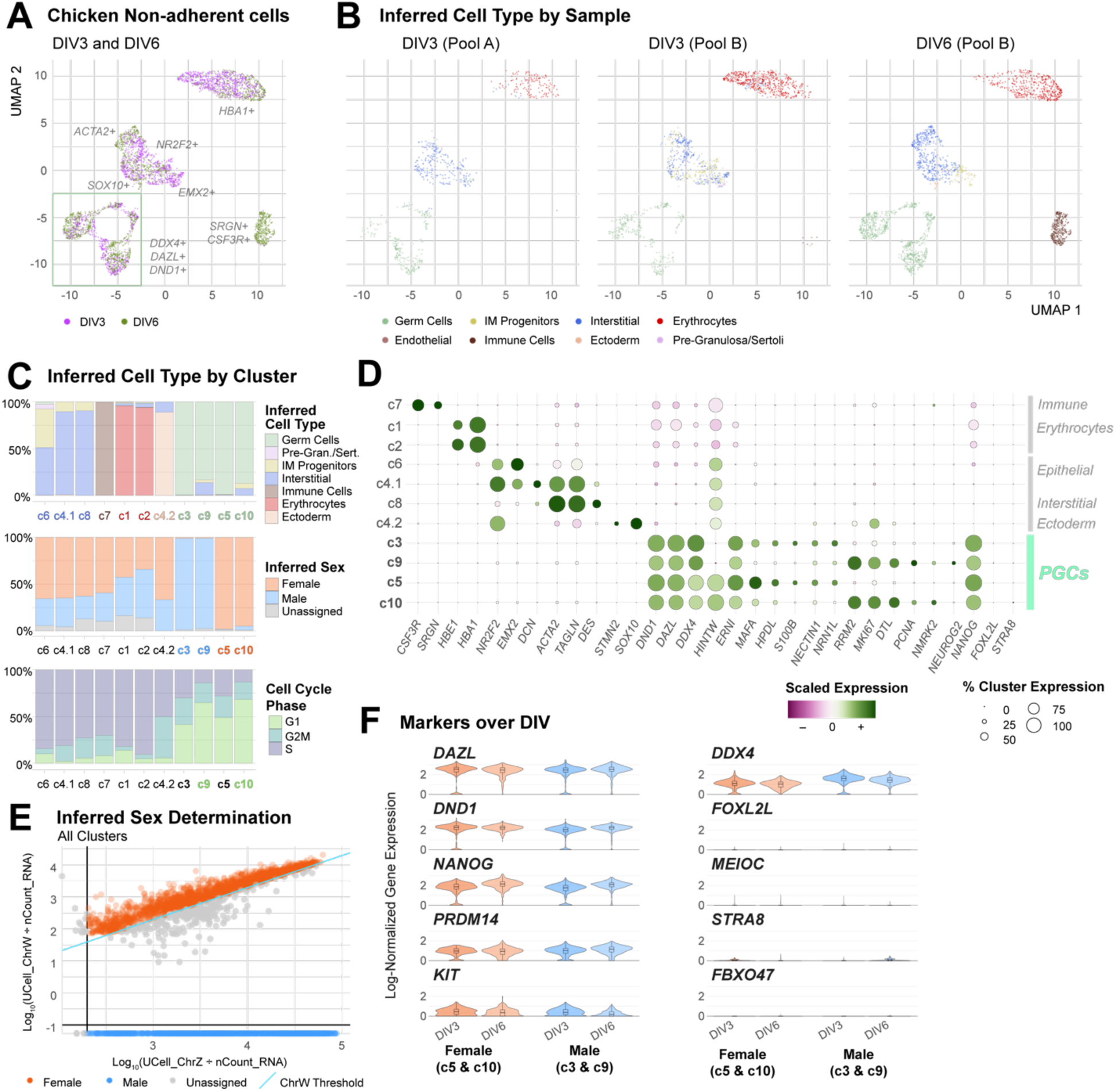
Processing and additional analysis for scRNAseq profiling of chicken short-term gonadal cell cultures. (**A**) UMAP dimensional reduction of integrated non-adherent chicken gonadal samples, colored by DIV and labeled by nearest-neighbor cluster assignment. Example cell type marker genes enriched in each cluster are indicated. (**B**) Same UMAP profile colored by inferred cell types at DIV3 and DIV6, and the different pools. (**C**) Proportional bar charts of inferred cell type, sex, or cell cycle phase by cluster. (**D**) Dotplot of select gene markers expressed in each cluster, highlighting cell type markers. (**E**) Scatterplot of log-normalized Chr.Z (x-axis) and Chr.W (y-axis) geneset expression, split by cluster. Dots denote geneset enrichments for each cell barcode, colored by inferred sex; line denotes a conservative inference threshold for barcode sex determination. (**F**) Violin Plot of select marker genes in germ cell clusters, split by inferred sex and DIV.

**Figure S6.**
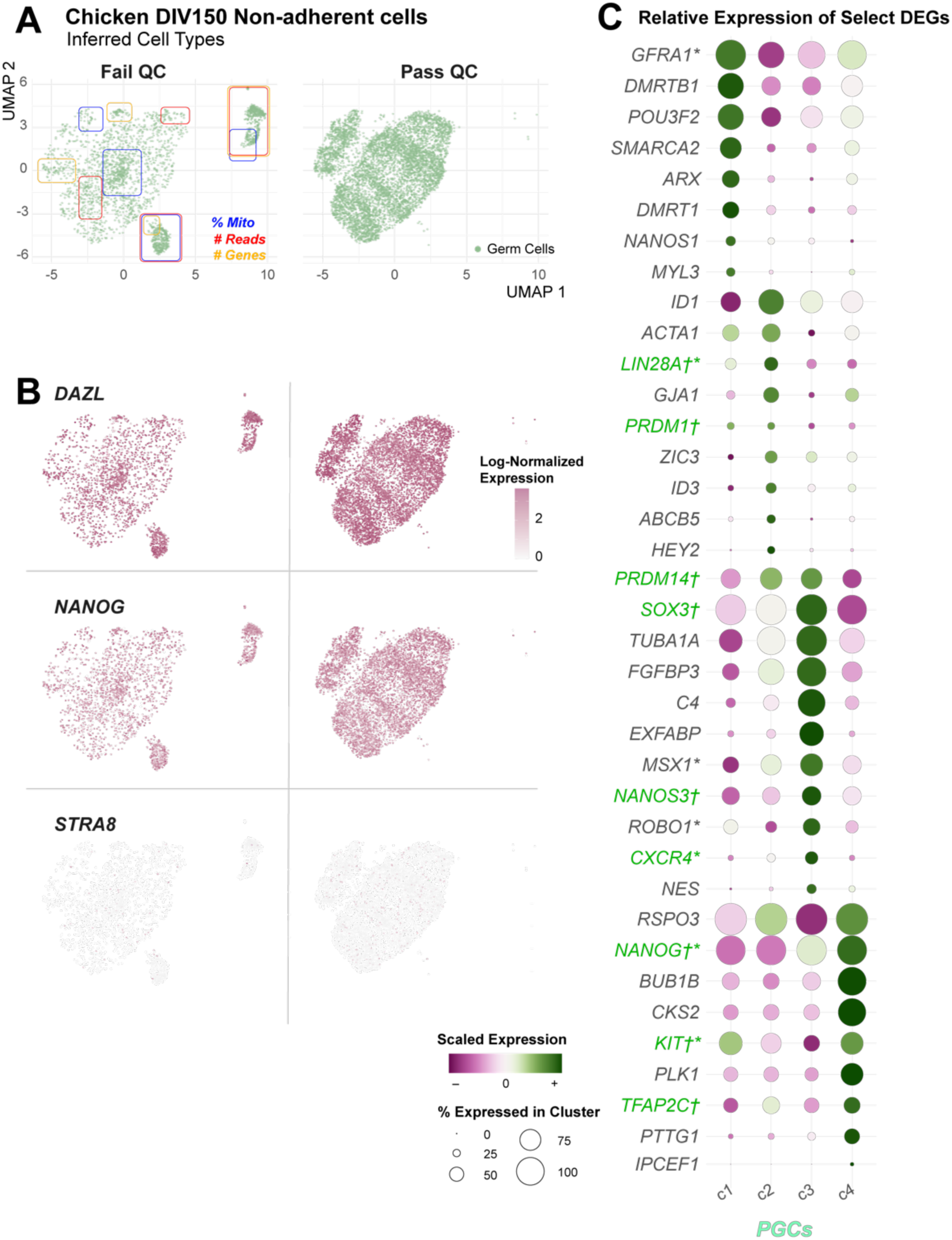
scRNAseq quality control and analysis of the DIV150 male chicken PGC cell line. (**A**) UMAP of minimally filtered dataset that failed or passed QC-based trimming, colored by inferred cell type (all cells were designated as germ cells). (**B**) Select germ cell gene expression markers overlaid onto the pass and failed cell datasets. (**C**) Dotplot of scaled gene expression between each cluster, highlighting select DEGs and canonical PGC cell type markers (green text). Genes are organized by cluster expression. * = genes associated with cell migration. † = genes that did not meet significant DEG thresholding (Log2FC ≥ -0.25; cluster expression ≥ 10%; adjusted p-value ≤ 0.05).

**Figure S7.**
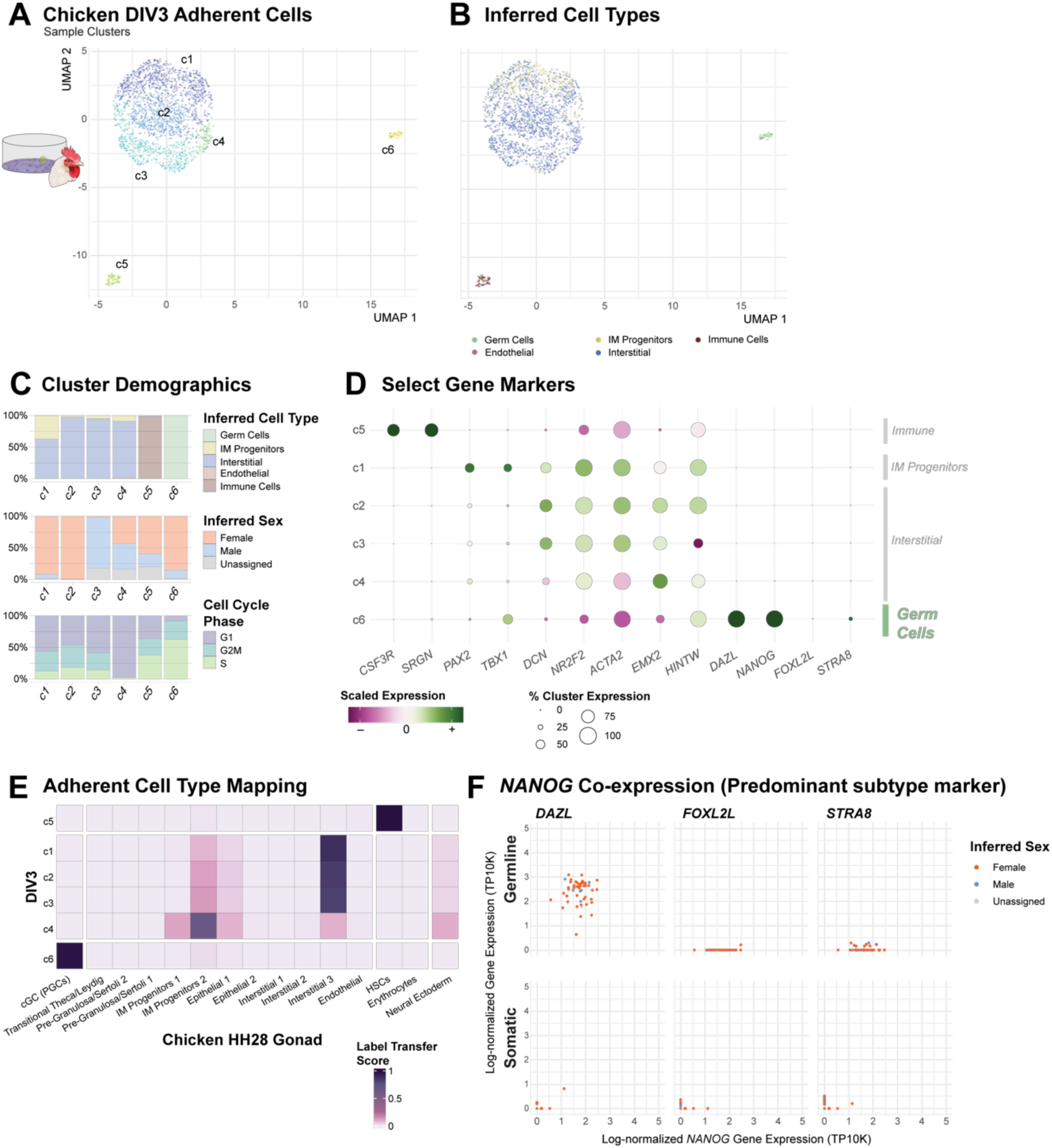
scRNAseq profiling of adherent cell types from DIV3 chicken gonadal cultures. (**A**) UMAP dimensional reduction of DIV3 adherent chicken gonadal samples, colored and labeled by nearest-neighbor cluster assignment. (**B**) The same UMAP, colored by inferred cell type. (**C**) Proportional bar charts of inferred cell type, sex, or cell cycle phase by cluster. (**D**) Dotplot of select gene markers expressed in each cluster, highlighting cell types. (**E**) Confusion matrix of cell type similarities between the chicken adherent cultured cells and the chicken *in vivo* HH28 gonadal dataset. (**F**) Co-expression analysis of germ cells (cluster c6) and somatic cells (all other clusters) from the adherent cells, colored by inferred sex. The dominant germ cell type marker, *NANOG,* is overlaid with other markers of differentiating chicken germ cells as determined by Biegler et al., 2024. The *STRA8* pre-meiotic marker is co-expressed with *NANOG* (PGC marker) as well as the lack of *FOXL2L* expression suggests these germ cell populations are not differentiating.

**Figure S8.**
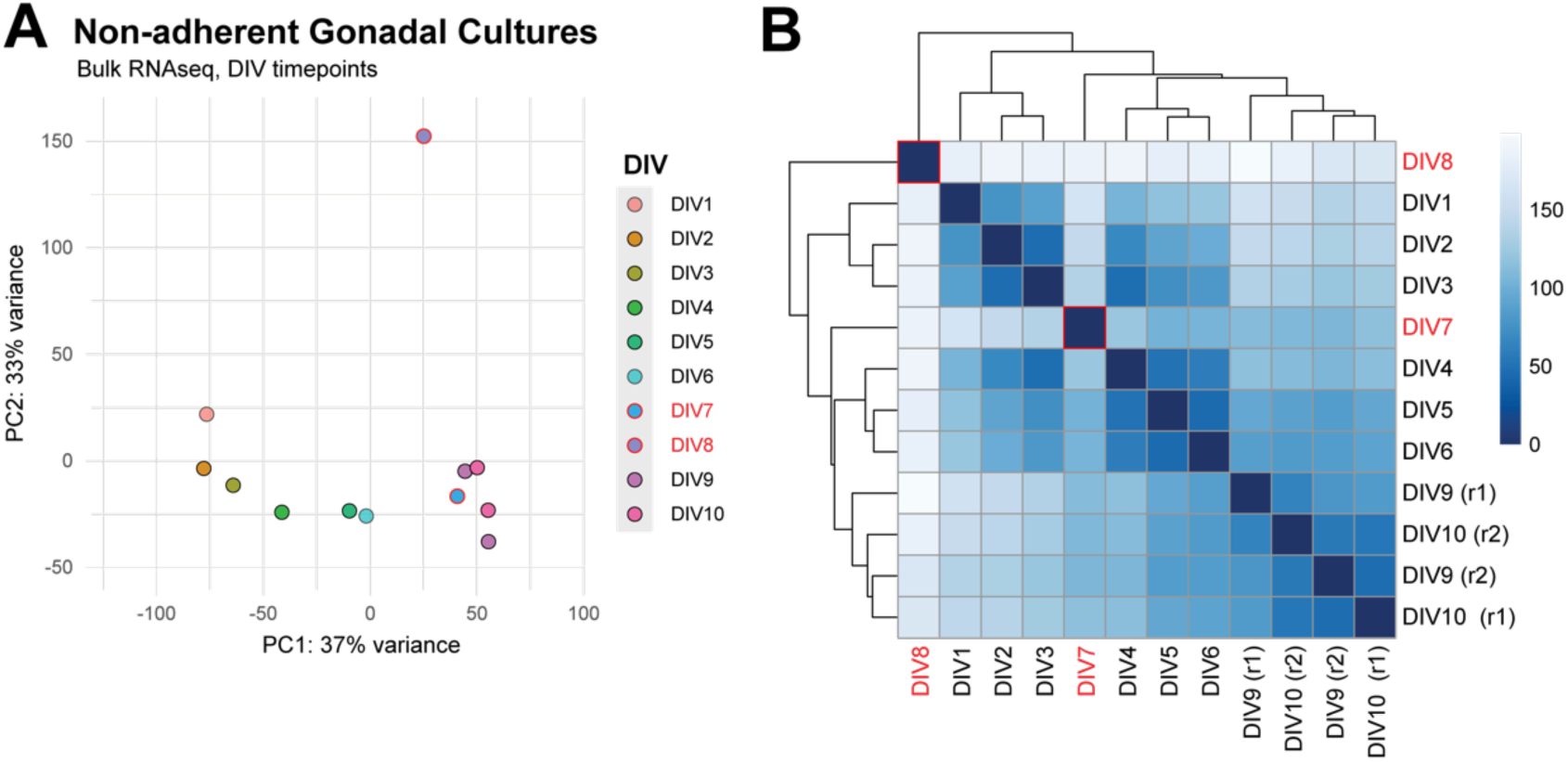
Bulk RNAseq quality control of gonadal cell culture time course. (**A**) PCA plot of DIV1 through DIV10 non-adherent culture transcriptome profiles. DIV7 and DIV8 are outliers. (**B**) Euclidean sample distance clustering between samples. Greater similarity (smaller Euclidean distance) is denoted by darker heatmap color. Note that DIV7 and DIV8 appear to cluster as outliers.

**Figure S9.**
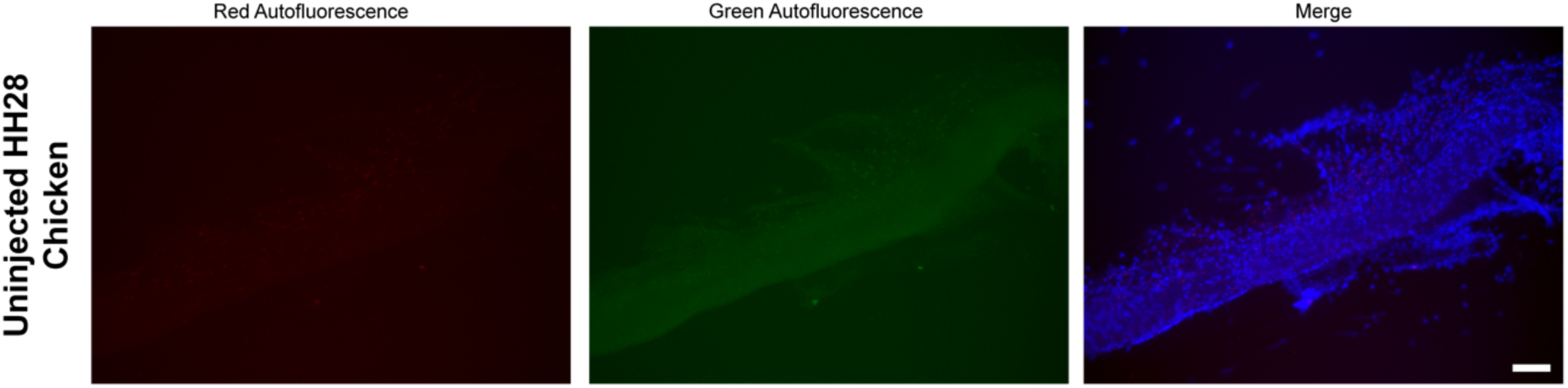
Exemplar single-channel images of dissected gonads from an uninjected ED6.5 chicken embryo. Blue labels DAPI nuclear stain. Scale bar = 50µm.

**Figure S10.**
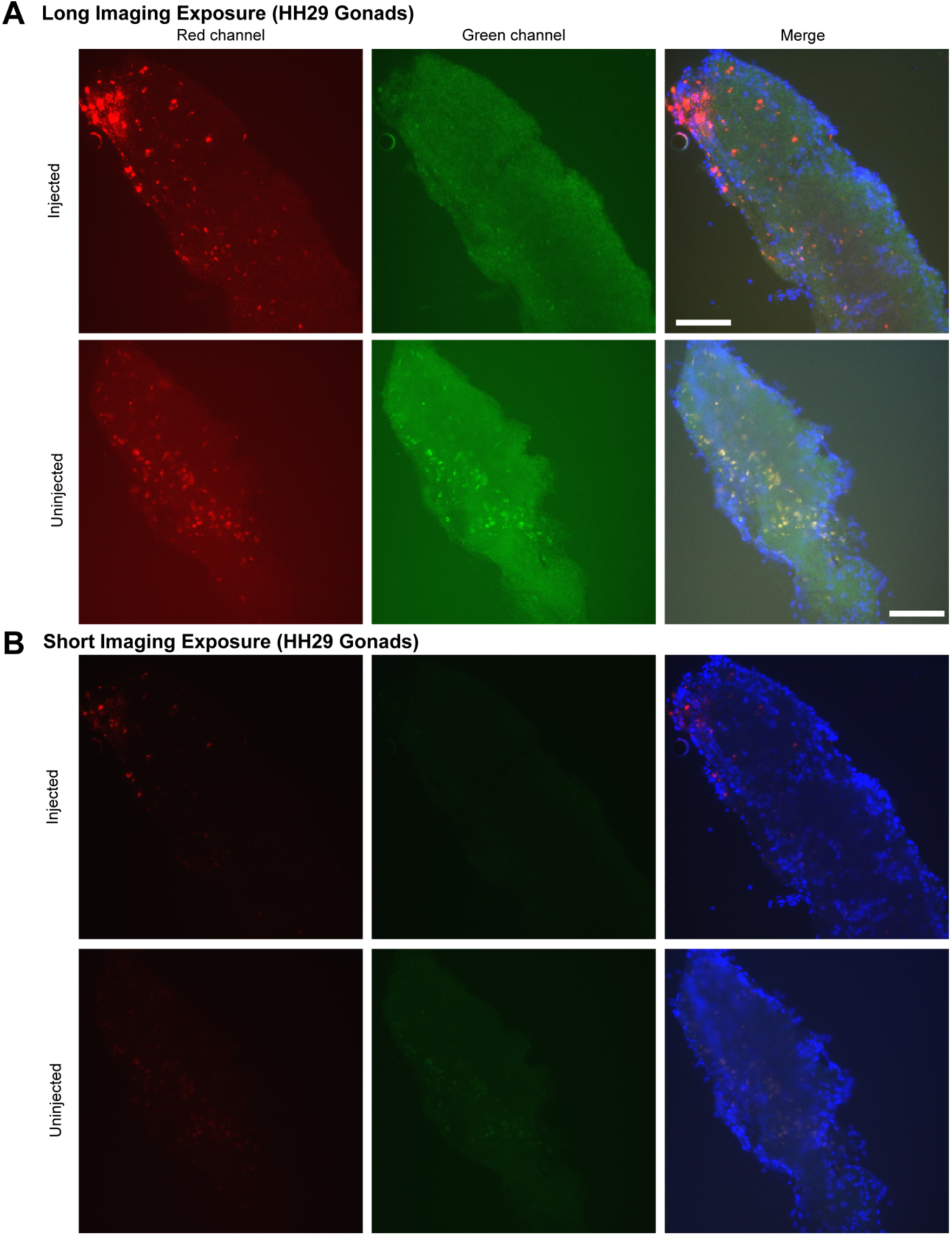
Assessment of PKH26 lipid membrane dye to assess the migratory competence of zebra finch gonadal cultures. (**A**) Long exposure images of dissected gonad from an ED6.5 zebra finch embryo injected with PKH26+ (red) non-adherent zebra finch gonadal cultures (top) or uninjected control ED6.5 zebra finch gonads (bottom). Note the autofluorescence (red and green) in the uninjected control. (**B**) Short exposure images of the same gonads. Note that while the uninjected control shows little autofluorescence, the injected gonad demonstrates inconsistent putative labeling that cannot be as reliably detected. Scale bars = 50µm.

**Figure S11.**
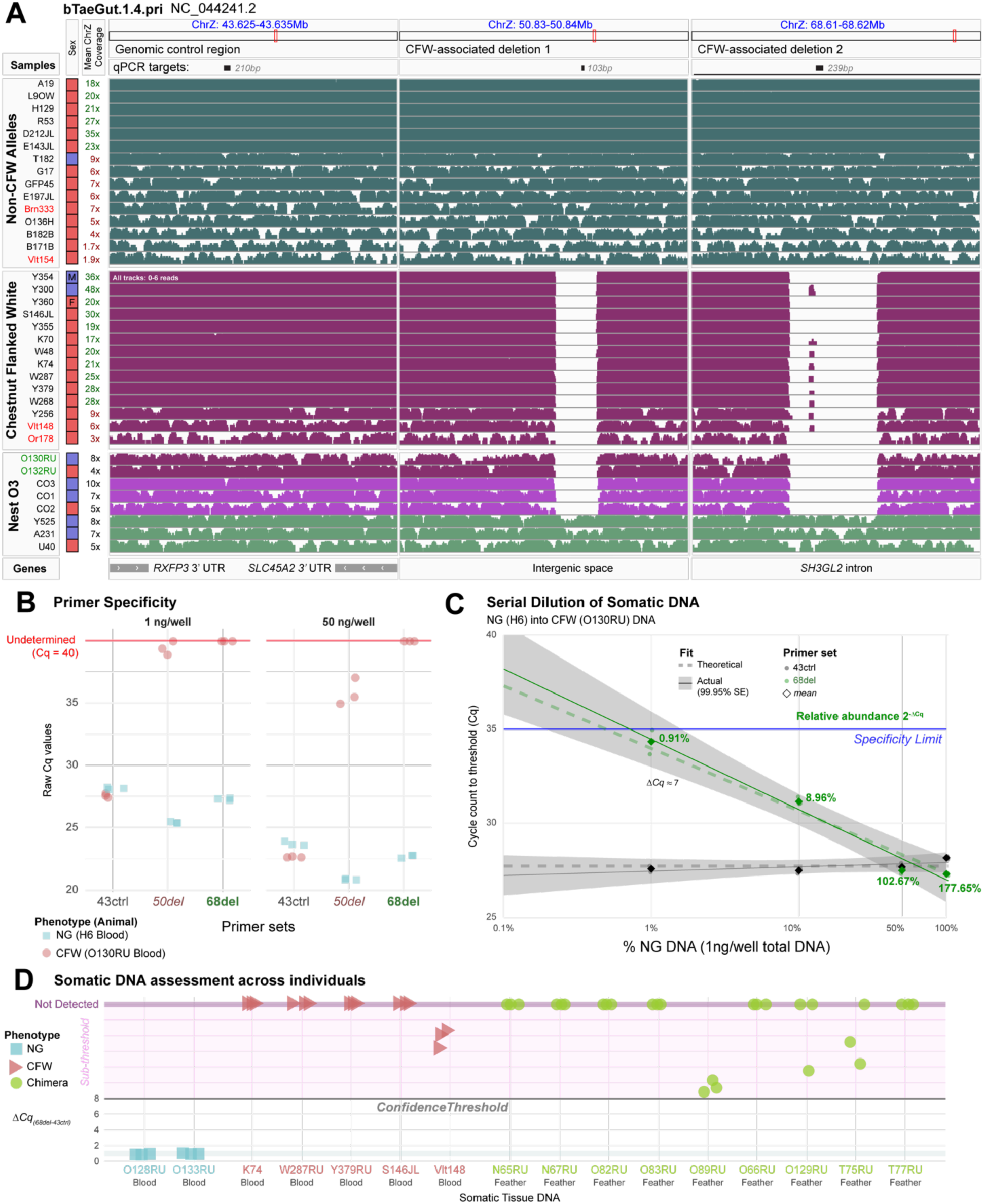
Identification of two deletions on Chr Z associated with the chestnut-flanked white (CFW) phenotype in the zebra finch. (**A**) IGV screenshot of natural grey (NG; n=15, top) and CFW (n=14, middle) short-read whole genome sequencing alignments, highlighting control (left) and CFW-deleted (middle and right) chromosome Z loci. Primer-targeted qPCR sites are annotated on top as black bars. Animal ID, sex and sequence depth are provided to on the left. Red labeled animal IDs are from non-Rockefeller populations. Alignments of nest O3 parents (green) and select offspring are also listed (bottom). (**B**) Genomic qPCR Cq values of the ChrZ primers using low (1ng/µL) and high (50ng/µL) DNA concentrations, assessing NG (blue square) and chimera (O130RU, red circle) DNA samples from blood. (**C**) The same blood DNA samples from B, titrated to simulate donor DNA using the 68del primer set. A specificity threshold (blue line) was estimated at Ct ≥ 35 based on the standard deviation of sample replicates at each titer. Percentages in green represent the relative abundance of 68del compared to 43ctrl primer sets, with the specificity limit reaching around 1% (ΔCq φ 7) for a 1ng/µL sample. (**D**) Validation of 68del qPCR on blood or feather DNA from multiple NG (blue squares), CFW (red triangle), or germline chimera (green circle) individuals. Note the low signal (ΔCq ≥ 8) in select CFW and chimera individuals, establishing a secondary confidence threshold below the specificity limit.

## 12 Supplemental Tables

**Table S1**. scRNAseq samples used for this study.

**Table S2**. Cell Barcode metadata for all scRNAseq objects.

**Table S3**. DEGs for each cluster in the zebra finch DIV3-6 non-adherent dataset.

**Table S4**. DEGs between zGC2 clusters (c1 and c2) in the zebra finch DIV3-6 non-adherent dataset.

**Table S5.** DEGs for each cluster in the zebra finch DIV3-6 adherent dataset.

**Table S6**. DEGs for each cluster in the chicken DIV3-6 non-adherent dataset.

**Table S7**. DEGs between the germ cell clusters in the chicken DIV3-6 non-adherent dataset.

**Table S8**. DEGs for each cluster in the chicken DIV150 non-adherent dataset.

**Table S9.** DEGs for each cluster in the chicken DIV3 adherent dataset.

**Table S10**. Bulk RNAseq samples used for this study.

**Table S11**. VST counts for Bulk RNAseq samples

**Table S12**. Exclusive cluster marker genesets (Table S3) used for ssGSEA to identify each zebra finch non-adherent cell types in bulk RNAseq data.

**Table S13**. Information for the egg injections for the gonadal migration assays.

**Table S14.** Zebra finch WGS sequencing samples used in this study.

**Table S15.** Cell culture media recipes used in this study.

**Table S16.** Primers used in this study.

## References

1. Petitte, J. N. Avian Germplasm Preservation: Embryonic Stem Cells or Primordial Germ Cells? Poultry Sci 85, 237–242 (2006).

2. Sun, Y., Li, Y., Zong, Y., Mehaisen, G. M. K. & Chen, J. Poultry genetic heritage cryopreservation and reconstruction: advancement and future challenges. J Anim Sci Biotechno 13, 115 (2022).

3. Swift, C. H. Origin and early history of the primordial germ-cells in the chick. Am. J. Anat. 15, 483– 516 (1914).

4. Bernardo, A. D. M., Sprenkels, K., Rodrigues, G., Noce, T. & Lopes, S. M. C. D. S. Chicken primordial germ cells use the anterior vitelline veins to enter the embryonic circulation. Biol Open 1, 1146–1152 (2012).

5. Park, T. S. & Han, J. Y. Conservation of Migration and Differentiation Circuits in Primordial Germ Cells Between Avian Species. J Reprod Develop 59, 252–257 (2013).

6. Whyte, J., Glover, J. D., Woodcock, M., Brzeszczynska, J., Taylor, L., Sherman, A., Kaiser, P. & McGrew, M. J. FGF, Insulin, and SMAD Signaling Cooperate for Avian Primordial Germ Cell Self-Renewal. Stem Cell Rep 5, 1171–1182 (2015).

7. Naito, M., Harumi, T. & Kuwana, T. Long Term in vitro Culture of Chicken Primordial Germ Cells Isolated from Embryonic Blood and Incorporation into Germline of Recipient Embryo. *J*. Poult. Sci. 47, 57 (2010).

8. Lavoir, M.-C. van de, Diamond, J. H., Leighton, P. A., Mather-Love, C., Heyer, B. S., Bradshaw, R., Kerchner, A., Hooi, L. T., Gessaro, T. M., Swanberg, S. E., Delany, M. E. & Etches, R. J. Germline transmission of genetically modified primordial germ cells. Nature 441, 766–769 (2006).

9. Choi, J. W., Kim, S., Kim, T. M., Kim, Y. M., Seo, H. W., Park, T. S., Jeong, J.-W., Song, G. & Han, J. Y. Basic Fibroblast Growth Factor Activates MEK/ERK Cell Signaling Pathway and Stimulates the Proliferation of Chicken Primordial Germ Cells. PloS one 5, e12968 (2010).

10. Park, T. S. & Han, J. Y. Derivation and characterization of pluripotent embryonic germ cells in chicken. Mol Reprod Dev 56, 475–482 (2000).

11. Kong, L., Qiu, L., Guo, Q., Chen, Y., Zhang, X., Chen, B., Zhang, Y. & Chang, G. Long-term in vitro culture and preliminary establishment of chicken primordial germ cell lines. Plos One 13, e0196459 (2018).

12. Tonus, C., Cloquette, K., Ectors, F., Piret, J., Gillet, L., Antoine, N., Desmecht, D., Vanderplasschen, A., Waroux, O. & Grobet, L. Long term-cultured and cryopreserved primordial germ cells from various chicken breeds retain high proliferative potential and gonadal colonisation competency. Reprod., Fertil. Dev. 28, 628–639 (2014).

13. Hu, T., Taylor, L., Sherman, A., Tiambo, C. K., Kemp, S. J., Whitelaw, B., Hawken, R. J., Djikeng, A. & McGrew, M. J. A low-tech, cost-effective and efficient method for safeguarding genetic diversity by direct cryopreservation of poultry embryonic reproductive cells. Elife 11, e74036 (2022).

14. Hu, T., Purdy, P. H., Blank, M. H., Muhonja, C. K., Pereira, R. J. G., Tiambo, C. K. & McGrew, M. J. Direct in vitro propagation of avian germ cells from an embryonic gonad biorepository. Poult. Sci. 104260 (2024). doi:10.1016/j.psj.2024.104260

15. Doddamani, D., Lázár, B., Ichikawa, K., Hu, T., Taylor, L., Gócza, E., Várkonyi, E. & McGrew, M. J. Propagation of goose primordial germ cells in vitro relies on FGF and BMP signalling pathways. Commun. Biol. 8, 301 (2025).

16. Park, K. J., Jung, K. M., Kim, Y. M., Lee, K. H. & Han, J. Y. Production of germline chimeric quails by transplantation of cryopreserved testicular cells into developing embryos. Theriogenology 156, 189– 195 (2020).

17. Chen, Y.-C., Lin, S.-P., Chang, Y.-Y., Chang, W.-P., Wei, L.-Y., Liu, H.-C., Huang, J.-F., Pain, B. & Wu, S.-C. In vitro culture and characterization of duck primordial germ cells. Poult. Sci. 98, 1820–1832 (2019).

18. Chaipipat, S., Prukudom, S., Sritabtim, K., Kuwana, T., Piyasanti, Y., Sinsiri, R., Piantham, C., Sangkalerd, S., Boonsanong, S., Pitiwong, K., Pidthong, A., Wanghongsa, S. & Siripattarapravat, K. Primordial germ cells isolated from individual embryos of red junglefowl and indigenous pheasants of Thailand. Theriogenology 165, 59–68 (2021).

19. Imus, N., Roe, M., Charter, S., Durrant, B. & Jensen, T. Transfer and Detection of Freshly Isolated or Cultured Chicken (Gallus gallus) and Exotic Species’ Embryonic Gonadal Germ Stem Cells in Host Embryos. Zool Sci 31, 360–368 (2014).

20. Borodin, P., Chen, A., Forstmeier, W., Fouché, S., Malinovskaya, L., Pei, Y., Reifová, R., Ruiz-Ruano, F. J., Schlebusch, S. A., Sotelo-Muñoz, M., Torgasheva, A., Vontzou, N. & Suh, A. Mendelian nightmares: the germline-restricted chromosome of songbirds. Chromosome Res 1–18 (2022). doi:10.1007/s10577-022-09688-3

21. Kinsella, C. M., Ruiz-Ruano, F. J., Dion-Côté, A.-M., Charles, A. J., Gossmann, T. I., Cabrero, J., Kappei, D., Hemmings, N., Simons, M. J. P., Camacho, J. P. M., Forstmeier, W. & Suh, A. Programmed DNA elimination of germline development genes in songbirds. Nature communications 10, 5468–10 (2019).

22. Katayama, M., Fukuda, T., Kaneko, T., Nakagawa, Y., Tajima, A., Naito, M., Ohmaki, H., Endo, D., Asano, M., Nagamine, T., Nakaya, Y., Saito, K., Watanabe, Y., Tani, T., Inoue-Murayama, M., Nakajima, N. & Onuma, M. Induced pluripotent stem cells of endangered avian species. Commun Biology 5, 1049 (2022).

23. Intarapat, S., Sukparangsi, W., Gusev, O. & Sheng, G. A Bird’s-Eye View of Endangered Species Conservation: Avian Genomics and Stem Cell Approaches for Green Peafowl (Pavo muticus). Genes 14, 2040 (2023).

24. Gessara, I., Dittrich, F., Hertel, M., Hildebrand, S., Pfeifer, A., Frankl-Vilches, C., McGrew, M. & Gahr, M. Highly Efficient Genome Modification of Cultured Primordial Germ Cells with Lentiviral Vectors to Generate Transgenic Songbirds. Stem Cell Rep (2021). doi:10.1016/j.stemcr.2021.02.015

25. Szczerba, A., Kuwana, T., Paradowska, M. & Bednarczyk, M. In Vitro Culture of Chicken Circulating and Gonadal Primordial Germ Cells on a Somatic Feeder Layer of Avian Origin. Animals 10, 1769 (2020).

26. Jung, K. M., Kim, Y. M., Keyte, A. L., Biegler, M. T., Rengaraj, D., Lee, H. J., Mello, C. V., Velho, T. A. F., Fedrigo, O., Haase, B., Jarvis, E. D. & Han, J. Y. Identification and characterization of primordial germ cells in a vocal learning Neoaves species, the zebra finch. FASEB J. 33, 13825–13836 (2019).

27. Jung, K. M., Kim, Y. M., Kim, J. L. & Han, J. Y. Efficient gene transfer into zebra finch germline-competent stem cells using an adenoviral vector system. Sci Rep-uk 11, 14746 (2021).

28. Jung, K. M., Kim, Y. M. & Han, J. Y. Transplantation and enrichment of busulfan-resistant primordial germ cells into adult testes for efficient production of germline chimeras in songbirds. Biol Reprod (2022). doi:10.1093/biolre/ioac206

29. Biegler, M. T., Belay, K., Wang, W., Szialta, C., Collier, P., Luo, J.-D., Haase, B., Gedman, G. L., Sidhu, A. V., Harter, E., Rivera-López, C., Amoako-Boadu, K., Fedrigo, O., Tilgner, H. U., Carroll, T., Jarvis, E. D. & Keyte, A. L. Pronounced early differentiation underlies zebra finch gonadal germ cell development. Dev. Biol. (2024). doi:10.1016/j.ydbio.2024.08.006

30. Ichikawa, K., Ezaki, R., Furusawa, S. & Horiuchi, H. Comparison of sex determination mechanism of germ cells between birds and fish: Cloning and expression analyses of chicken forkhead box L3-like gene. Dev. Dyn. 248, 826–836 (2019).

31. Smith, C. A., Roeszler, K. N., Bowles, J., Koopman, P. & Sinclair, A. H. Onset of meiosis in the chicken embryo; evidence of a role for retinoic acid. Bmc Dev Biol 8, 85–85 (2008).

32. Pigozzi, M. I. & Solari, A. J. Germ cell restriction and regular transmission of an accessory chromosome that mimics a sex body in the zebra finch, Taeniopygia guttata. Chromosome Res 6, 105– 113 (1998).

33. Torgasheva, A. A., Malinovskaya, L. P., Zadesenets, K. S., Karamysheva, T. V., Kizilova, E. A., Akberdina, E. A., Pristyazhnyuk, I. E., Shnaider, E. P., Volodkina, V. A., Saifitdinova, A. F., Galkina, S. A., Larkin, D. M., Rubtsov, N. B. & Borodin, P. M. Germline-restricted chromosome (GRC) is widespread among songbirds. Proceedings of the National Academy of Sciences 116, 11845–11850 (2019).

34. Smith, J. J., Timoshevskiy, V. A. & Saraceno, C. Programmed DNA Elimination in Vertebrates. Annu. Rev. Anim. Biosci. 9, 1–29 (2020).

35. Zou, X., He, Y., Zhao, Z., Li, J., Qu, H., Liu, Z., Chen, P., Ji, J., Zhao, H., Shu, D. & Luo, C. Single-cell RNA-seq offer new insights into the cell fate decision of the primordial germ cells. Int. J. Biol. Macromol. 139136 (2024). doi:10.1016/j.ijbiomac.2024.139136

36. Jung, K. M., Seo, M. & Han, J. Y. Comparative single-cell transcriptomic analysis reveals differences in signaling pathways in gonadal primordial germ cells between chicken (Gallus gallus) and zebra finch (Taeniopygia guttata). Faseb J 37, e22706 (2023).

37. Magnúsdóttir, E., Dietmann, S., Murakami, K., Günesdogan, U., Tang, F., Bao, S., Diamanti, E., Lao, K., Gottgens, B. & Surani, M. A. A tripartite transcription factor network regulates primordial germ cell specification in mice. Nat Cell Biol 15, 905–915 (2013).

38. Sánchez-Sánchez, A. V., Camp, E., Leal-Tassias, A., Atkinson, S. P., Armstrong, L., Díaz-Llopis, M. & Mullor, J. L. Nanog Regulates Primordial Germ Cell Migration Through Cxcr4b. Stem Cells 28, 1457– 1464 (2010).

39. Suzuki, K., Kwon, S. J., Saito, D. & Atsuta, Y. LIN28 is essential for the maintenance of chicken primordial germ cells. Cells Dev. 176, 203874 (2023).

40. Srihawong, T., Kuwana, T., Siripattarapravat, K. & Tirawattanawanich, C. Chicken primordial germ cell motility in response to stem cell factor sensing. Int. J. Dev. Biol. 59, 453–460 (2016).

41. Tomasz, M. Mitomycin C: small, fast and deadly (but very selective). Chem. Biol. 2, 575–579 (1995).

42. Gessara, I. Generation of transgenic zebra finches by the culture and genetic modification of germline stem cells. (2021).

43. Landry, G. P. An Almost Complete Guide to: The Varieties and Genetics of the Zebra Finch. (Poule d’eau Publishing Co., 1997). at <http://www.zebrafinch.com/other/Book.html≥

44. Gunnarsson, U., Hellström, A. R., Tixier-Boichard, M., Minvielle, F., Bed’hom, B., Ito, S., Jensen, P., Rattink, A., Vereijken, A. & Andersson, L. Mutations in SLC45A2 Cause Plumage Color Variation in Chicken and Japanese Quail. Genetics 175, 867–877 (2007).

45. Roy, S. G., Bakhrat, A., Abdu, M., Afonso, S., Pereira, P., Carneiro, M. & Abdu, U. Mutations in SLC45A2 lead to loss of melanin in parrot feathers. G3: Genes, Genomes, Genet. 14, jkad254 (2023).

46. Xu, X., Dong, G.-X., Hu, X.-S., Miao, L., Zhang, X.-L., Zhang, D.-L., Yang, H.-D., Zhang, T.-Y., Zou, Z.-T., Zhang, T.-T., Zhuang, Y., Bhak, J., Cho, Y. S., Dai, W.-T., Jiang, T.-J., Xie, C., Li, R. & Luo, S.-J. The genetic basis of white tigers. Curr Biology Cb 23, 1031–5 (2013).

47. Manichaikul, A., Mychaleckyj, J. C., Rich, S. S., Daly, K., Sale, M. & Chen, W.-M. Robust relationship inference in genome-wide association studies. Bioinformatics 26, 2867–2873 (2010).

48. Estermann, M. A., Williams, S., Hirst, C. E., Roly, Z. Y., Serralbo, O., Adhikari, D., Powell, D., Major, A. T. & Smith, C. A. Insights into Gonadal Sex Differentiation Provided by Single-Cell Transcriptomics in the Chicken Embryo. Cell Reports 31, 107491 (2020).

49. Estermann, M. A., Mariette, M. M., Moreau, J. L. M., Combes, A. N. & Smith, C. A. PAX2+ Mesenchymal Origin of Gonadal Supporting Cells Is Conserved in Birds. Front. Cell Dev. Biol. 9, 735203 (2021).

50. Biederman, M. K., Nelson, M. M., Asalone, K. C., Pedersen, A. L., Saldanha, C. J. & Bracht, J. R. Discovery of the First Germline-Restricted Gene by Subtractive Transcriptomic Analysis in the Zebra Finch, Taeniopygia guttata. Current biology : CB 28, 1620–1627.e5 (2018).

51. Liu, Y., Kossack, M. E., McFaul, M. E., Christensen, L. N., Siebert, S., Wyatt, S. R., Kamei, C. N., Horst, S., Arroyo, N., Drummond, I. A., Juliano, C. E. & Draper, B. W. Single-cell transcriptome reveals insights into the development and function of the zebrafish ovary. Elife 11, e76014 (2022).

52. Tanaka, M. Germline stem cells are critical for sexual fate decision of germ cells. Bioessays 38, 1227– 1233 (2016).

53. Ishiguro, K., Matsuura, K., Tani, N., Takeda, N., Usuki, S., Yamane, M., Sugimoto, M., Fujimura, S., Hosokawa, M., Chuma, S., Ko, M. S. H., Araki, K. & Niwa, H. MEIOSIN Directs the Switch from Mitosis to Meiosis in Mammalian Germ Cells. Dev Cell 52, 429–445.e10 (2020).

54. Zhang, Y., Wang, Y., Zuo, Q., Li, D., Zhang, W., Wang, F., Ji, Y., Jin, J., Lu, Z., Wang, M., Zhang, C. & Li, B. CRISPR/Cas9 mediated chicken Stra8 gene knockout and inhibition of male germ cell differentiation. Plos One 12, e0172207 (2017).

55. Oulad-Abdelghani, M., Bouillet, P., Décimo, D., Gansmuller, A., Heyberger, S., Dollé, P., Bronner, S., Lutz, Y. & Chambon, P. Characterization of a premeiotic germ cell-specific cytoplasmic protein encoded by Stra8, a novel retinoic acid-responsive gene. J. cell Biol. 135, 469–477 (1996).

56. Nakamura, Y. Avian Reproduction, From Behavior to Molecules. Adv. Exp. Med. Biol. 1001, 187– 214 (2017).

57. IUCN. The IUCN Red List of Threatened Species. (2022). at <https://www.iucnredlist.org≥

58. Agate, R. J., Scott, B. B., Haripal, B., Lois, C. & Nottebohm, F. Transgenic songbirds offer an opportunity to develop a genetic model for vocal learning. Proc National Acad Sci 106, 17963–17967 (2009).

59. Shvedov, N. R., Analoui, S., Dafalias, T., Bedell, B. L., Gardner, T. J. & Scott, B. B. In vivo imaging in transgenic songbirds reveals superdiffusive neuron migration in the adult brain. Cell Rep. 43, 113759 (2024).

60. Iikawa, H., Nishina, A., Morita, M., Atsuta, Y., Hayashi, Y. & Saito, D. Labeling and sorting of avian primordial germ cells utilizing Lycopersicon Esculentum lectin. *Dev.*, Growth Differ. (2024). doi:10.1111/dgd.12948

61. Jung, J. G., Kim, D. K., Park, T. S., Lee, S. D., Lim, J. M. & Han, J. Y. Development of Novel Markers for the Characterization of Chicken Primordial Germ Cells. Stem Cells 23, 689–698 (2005).

62. Kim, Y.-K., Cho, B., Cook, D. P., Trcka, D., Wrana, J. L. & Ramalho-Santos, M. Absolute scaling of single-cell transcriptomes identifies pervasive hypertranscription in adult stem and progenitor cells. Cell Reports 42, 111978 (2023).

63. Kim, Y.-K., Collignon, E., Martin, S. B. & Ramalho-Santos, M. Hypertranscription: the invisible hand in stem cell biology. Trends Genet. (2024). doi:10.1016/j.tig.2024.08.005

64. Langmead, B., Trapnell, C., Pop, M. & Salzberg, S. L. Ultrafast and memory-efficient alignment of short DNA sequences to the human genome. Genome Biol. 10, R25 (2009).

65. Picard Toolkit. Broad Institute, GitHub repository (2019). at <https://broadinstitute.github.io/picard/>

66. Robinson, J. T., Thorvaldsdóttir, H., Winckler, W., Guttman, M., Lander, E. S., Getz, G. & Mesirov, J. P. Integrative genomics viewer. Nat Biotechnol 29, 24–26 (2011).

67. Danecek, P., Auton, A., Abecasis, G., Albers, C. A., Banks, E., DePristo, M. A., Handsaker, R. E., Lunter, G., Marth, G. T., Sherry, S. T., McVean, G., Durbin, R. & Group, 1000 Genomes Project Analysis. The variant call format and VCFtools. Bioinformatics 27, 2156–2158 (2011).

68. Kucera, A. C. & Heidinger, B. J. Avian Semen Collection by Cloacal Massage and Isolation of DNA from Sperm. J Vis Exp (2018). doi:10.3791/55324

69. Ye, J., Coulouris, G., Zaretskaya, I., Cutcutache, I., Rozen, S. & Madden, T. L. Primer-BLAST: A tool to design target-specific primers for polymerase chain reaction. BMC Bioinform. 13, 134 (2012).

70. Wallace, E. & Haynes, S. The tidyqpcr R Package: Quantitative PCR analysis in the tidyverse. (2021). at https://github.com/ewallace/tidyqpcr

